# Decoding conformational heterogeneity across disordered proteomes

**DOI:** 10.64898/2026.03.13.711260

**Authors:** Anton Abyzov, Markus Zweckstetter

**Affiliations:** German Center for Neurodegenerative Diseases (DZNE), Von-Siebold-Str. 3a, 37075 Göttingen, Germany; brainQr Therapeutics GmbH, Von-Siebold-Str. 3a, 37075 Göttingen, Germany

## Abstract

Intrinsically disordered proteins (IDPs) comprise nearly one-third of the human proteome and play key roles in regulation, signaling, and disease, yet their dynamic nature has resisted accurate structural prediction even by the state of the art deep-learning methods. Here we introduce AI-IDP, a framework combining established deep-learning structure prediction of isolated short IDP fragments with their flexible physical restrains-aware assembly to transform sequence information into experiment-consistent conformational ensembles of disordered proteins. AI-IDP reproduces experimental observables across local, medium-range, and global scales, including transient secondary structure, mutation sensitivity, and overall chain dimensions. Applied to more than 3,000 disordered regions across human and non-human proteomes, AI-IDP reveals that transient α-helices and polyproline-II conformations are pervasive and evolutionarily tuned features of disorder. By uncovering how sequence encodes conformational heterogeneity, AI-IDP provides a practical framework for understanding the structural and functional logic of disordered proteomes and enables rationally targeting the dynamic protein states that underlie health and disease.

## Main

Proteins without a fixed three-dimensional structure – intrinsically disordered proteins (IDPs) – represent one of biology’s most abundant and enigmatic classes ^1,2^. Comprising nearly one-third of the human proteome, they regulate essential signaling, transcriptional, and stress-response pathways, and are strongly implicated in neurodegenerative and malignant diseases ^1,3^. Yet, because they lack a stable fold, IDPs challenge the classical structure-function paradigm that has defined molecular biology for decades. Their conformational landscapes are dynamic, heterogeneous, and only partially structured, and these fleeting elements of order are critical for recognition, regulation, and phase behavior ^1–3^.

Understanding how sequence encodes such dynamic behavior remains a major challenge. Despite their importance, no existing sequence-based method can consistently predict conformational ensembles of IDPs that accurately capture their structural features across scales with experimental concordance ^4,5^. Long-timescale molecular dynamics simulations oversample structure ^6,7^, produce spurions intramolecular contacts (ref. ^6^ SI Fig. S5-S8) and often require extended computational times and specialized hardware. Established deep-learning predictors such as AlphaFold2, which have transformed folded-protein prediction,^8^ often fail to capture flexibility and produce spuriously ordered states ^9^, an issue that reappears in some emerging AI-driven IDP ensemble predictors. Coarse grained models^10–12^, on the other hand, often miss transiently folded elements in generated IDP ensembles. Thus, in many cases local conformational preferences, such as secondary structure propensities, and global conformational properties, such as hydrodynamic radius, are often tuned in parallel.

Here we introduce AI-IDP, a framework that generates experiment-consistent all-atom conformational ensembles of IDPs directly from sequence. Instead of developing a disordered conformation predictor for an entire IDP, we use deep-learning models developed for folded proteins (AlphaFold2) to generate conformations of short overlapping IDP fragments. Combined with a flexible assembly of fragment structures that takes into account physical restraints (backbone angles and steric clashes), AI-IDP bridges the gap between sequence-based AI models and the predominantly local nature of transient structure in IDPs. We validate the approach across diverse IDPs, reveal sequence-dependent transient structure and mutation sensitivity, and extend ensemble prediction to thousands of disordered regions across proteomes. The results establish a framework linking sequence to structural heterogeneity and reveal how evolution shapes the conformational logic of disorder in living systems.

### Capturing flexibility in intrinsically disordered proteins

The dynamic nature of IDPs arises from their ability to sample a continuum of conformations rather than a single folded structure. To capture this intrinsic flexibility, we designed AI-IDP. AI-IDP represents an IDP chain as a series of 10-residue overlapping fragments whose local conformations are predicted by the established deep-learning protein structure model AlphaFold2. To grow the full IDP chain from short overlapping fragments, the two last residues of a precedent fragment were overlapped with the first two residues of a subsequent fragment. Subsequently, the backbone dihedral angles in the linking region were resampled using a library of amino acid-specific values extracted from loop regions of folded proteins^13^ to keep IDP chains more flexible while also avoiding steric clashes. Dihedral angles in the linking region, however, were not changed when a helical structure was present in order to retain transient helical structure. Conformational variation was further introduced through random selection of fragments from the five AlphaFold2-predicted structures. This combined approach allowed local structural preferences to emerge from sequence while preserving the global disorder of the chain (Extended Data Fig. 1). Further details of the approach are presented in the Methods section (see also Extended Data Figs. 1-7; Extended Data Table 1). AI-IDP thus generates a large set of all-atom conformers that collectively define an ensemble distribution. These ensembles exhibit high structural diversity and lack stable tertiary contacts, consistent with experimental descriptions of disorder. We then asked whether this fragment-based representation can recover transient local structure, mutation effects, medium-range contacts and overall chain dimensions in well-characterized IDPs (Extended Data Table 2).

### Sequence – structure relationships

We next tested whether AI-IDP can recover transient local structure in intrinsically disordered proteins directly from sequence. For six benchmark IDPs – c-Myc, ACTR, 4E-BP2, α-synuclein, Tau, and p53 – AI-IDP ensembles were compared with experimental NMR data that define their transient secondary-structure propensities (Extended Data Table 2).

Across all six proteins, AI-IDP accurately reproduced residue-specific secondary structure profiles derived from experimental C_α_/C_β_ chemical shifts (see Methods), with correlation coefficients up to 0.9 and root-mean-square deviations below 0.15 (Fig. 1A (ii-vii); Extended Data Fig. 8). Raw C_α_/C_β_ chemical shifts are also reproduced by AI-IDP with high accuracy (Extended Data Fig. 9). The predicted ensembles captured α-helical and extended segments observed experimentally, while avoiding the over-stabilized helices generated by AlphaFold2. For example, AI-IDP recovered the transient helix in the c-Myc region that mediates binding to Max, and the dynamic helices in ACTR that form stable structure upon coactivator binding - features absent from static prediction models.

**Figure 1.**
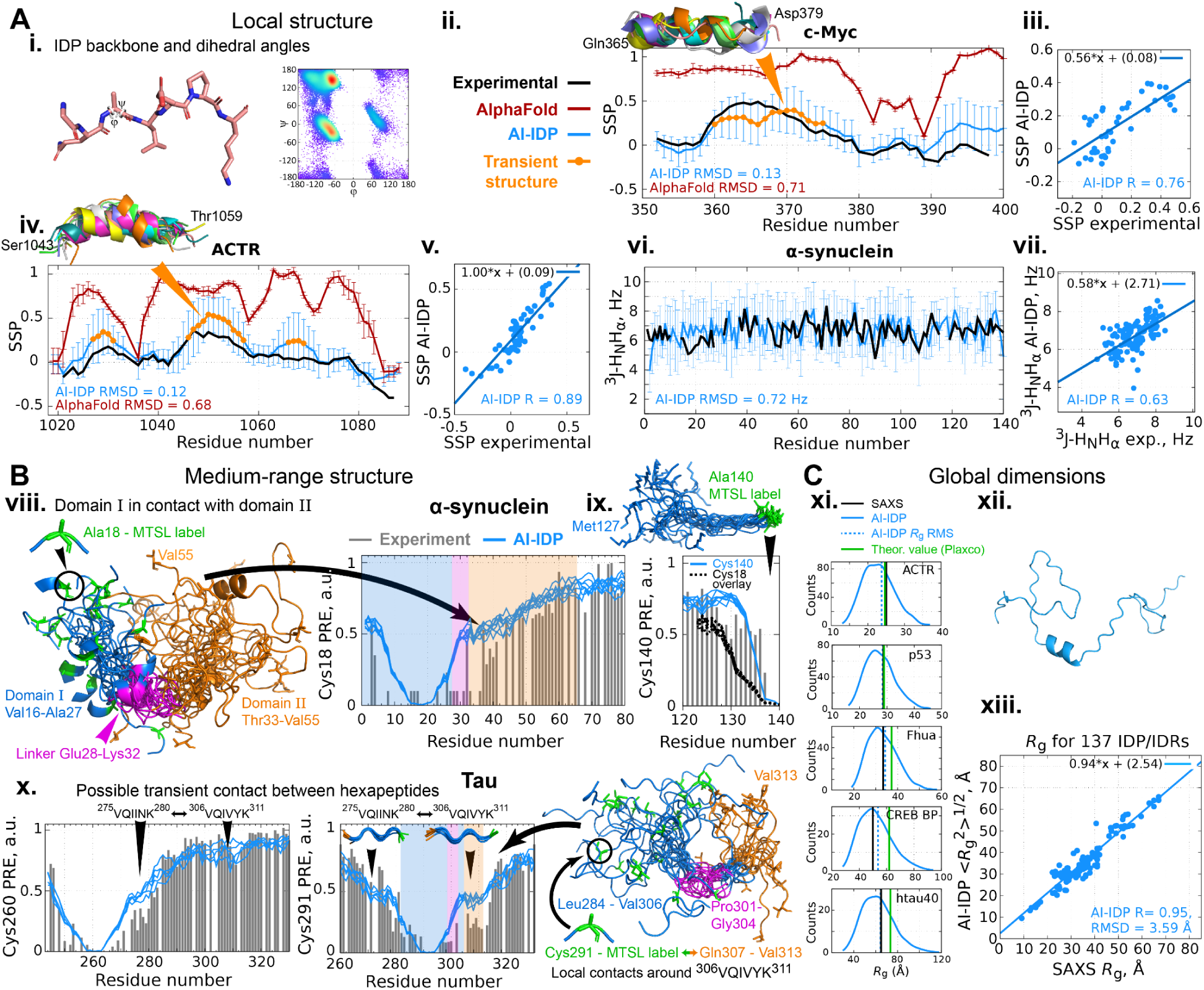
Decoding conformational heterogeneity in disordered proteins from local to global scale. **A.** Local structural propensities recovered by AI-IDP. (i) Representative fragment from an AI-IDP ensemble and corresponding Ramachandran distribution of backbone ϕ, ψ angles, illustrating conformational diversity. (ii-iii) Residue-specific SSP scores for c-Myc obtained from experimental C_α_/C_β_ NMR chemical shifts (black), AI-IDP ensembles (blue), and AlphaFold2 predictions (brown). Regions with significant transient structure (SSP > 0.2 for ≥ 3 consecutive residues) are highlighted in orange. (iv-v) Equivalent comparison for ACTR, showing quantitative agreement between AI-IDP and experiment. (vi-vii) Experimental versus AI-IDP-predicted ^3^J(H_N_H_α_) couplings for α-synuclein demonstrate accurate reproduction of local backbone flexibility. **B,** Medium-range contacts revealed by paramagnetic relaxation enhancement (PRE). (viii) N-terminal region of the α-synuclein AI-IDP ensemble showing transient contact between domains I and II; experimental (black) and AI-IDP-predicted (blue) PRE profiles for α-synuclein with a spin label at Cys18. (ix) Experimental and AI-IDP-predicted PRE profiles for α-synuclein with a spin label at Cys140; black dashed line shows the Cys18 profile for comparison. (x) Experimental and AI-IDP-predicted PREs for Tau K18 with spin labels at Cys260 or Cys291, together with representative ensemble conformers. Five 1000-residue AI-IDP ensembles of α-synuclein and Tau K18 were used for PRE predictions, shown as five distinct blue lines, to demonstrate structural features reproducibility. **C,** Global chain dimensions captured by AI-IDP. (xi) Distributions of radii of gyration (*R*_g_) from AI-IDP ensembles for five IDPs, compared with SAXS-derived and theoretical random-coil values ^33^. (xii) Representative AI-IDP conformer of ACTR. (xiii) Correlation between SAXS-derived *R*_g_ and ensemble-averaged *R*_g_ for 137 IDPs (r = 0.95, RMSD = 3.6 Å). Together, the analyses demonstrate that AI-IDP reproduces experimental observables from local secondary-structure propensities to global compaction, capturing the hierarchical organization of disordered protein ensembles.

The ability to model intrinsic flexibility was particularly evident for the 140-residue α-synuclein, whose AI-IDP ensemble displays heterogeneous conformations with a partially helical N-terminus, in agreement with NMR-derived SSP and ^3^J-H_N_H_α_ coupling data (Fig. 1A (vi-vii) and Extended Data Fig. 10). When flexibility is not incorporated, AlphaFold2 collapses α-synuclein into a single extended α-helix. Similarly, AI-IDP captured transient helices at the termini of Tau and within the N-terminal transactivation domain of p53, consistent with experimental observations that these elements mediate partner recognition.

Comparisons with other ensemble generators further emphasize AI-IDP’s accuracy. We used experimental chemical shifts for 40 different IDP/IDRs (Extended Data Table 3). CALVADOS ^14^ and Flexible-Meccano ^15^ fail to reproduce short transient helices in IDPs, resulting in lower correlations with experiment and higher RMSD values (Figs. 2A, 2C; Extended Data Figs. 11, 12). idpGAN ^11^ and STARLING ^12^ also demonstrate low correlations with experiment and higher RMSD values. IDPFold ^16^, IDPFold2 ^17^ and IDPConformerGenerator ^18^ demonstrate higher correlations with experiment (Fig. 2A). However, there is a high amount of spuriously predicted transient structure in these models (Fig. 2D, Extended Data Figs. 13, 14), resulting in increased RMSDs with experimental data. In addition, a recently presented ‘disorder-specialized predictor’ IDP-o ^19^ that utilizes assemblies of fragments derived from structures in the AlphaFold Protein Structure Database also demonstrates higher RMSDs with experiment and a significant number of residues with overpredicted transient structure (Fig. 2D). This underscores the importance of a *de novo* structure prediction of isolated IDP fragments, employed in AI-IDP, rather than fragment extraction from existing AlphaFold structures.

**Figure 2.**
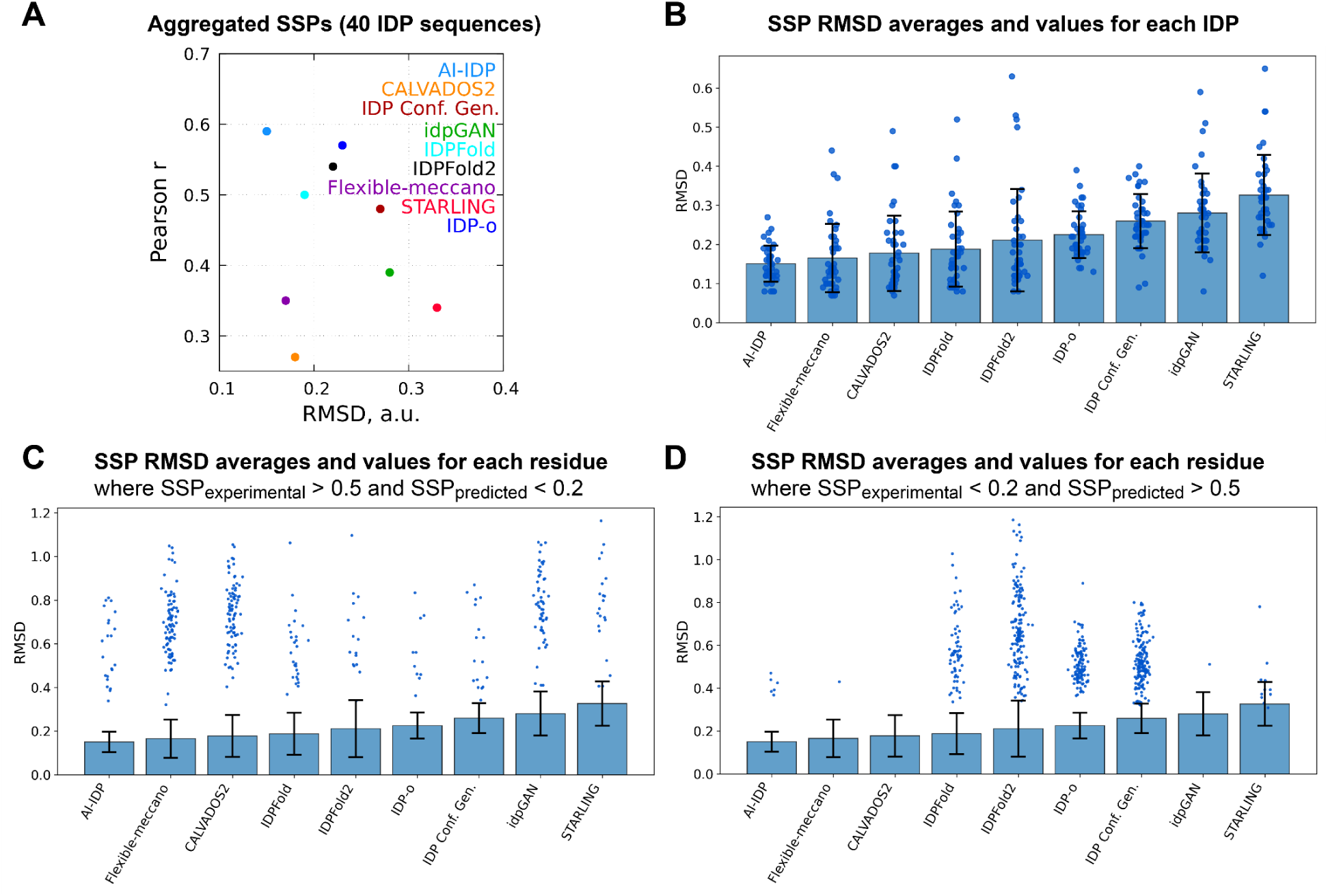
Sequence-encoded local conformational biases are recovered by AI-IDP with high ac-curacy. **A.** Accuracy of local structural properties in AI-IDP (light blue), CALVADOS2 (orange), IDP Conformer Generator (red-brown), idpGan (green), IDPFold (cyan), IDPFold2 (black), Flexible-Meccano (purple), STARLING (magenta) and IDP-o (blue) ensembles. Each dot represents a comparison between ensemble-derived and experimental SSPs for aggregated SSPs of 40 IDPs (Extended Data Table 3). The horizontal and vertical axes represent RMSDs and Pearson correlation coefficients between ensemble-derived and experimental SSPs, respectively. Parameters used for ensemble generation are specified in the Methods section. AI-IDP achieves highest SSP correlation values and lowest RMSDs. **B.** Accuracy of local structural properties as measured by IDP-specific and average RMSDs between ensemble-derived and experimental SSPs. AI-IDP achieves lowest average RMSDs with no IDPs exceeding RMSD value of 0.3. **C, D.** Accuracy of residue-level SSP predictions by different methods. The dot spread above blue bars represent RMSD values for residues where experimentally-observed transient structure is significantly underestimated^11^ (C; experimental SSP > 0.5 and predicted SSP < 0.2) or, inversely, overestimated (D; experimental SSP < 0.2 and predicted SSP < 0.5). Overall, only AI-IDP provides accurate estimates of secondary structure in sampled IDPs.

To further benchmark AI-IDP against physics-based simulations and assess its computational efficiency, we compared ensembles generated by AI-IDP with those from extensive all-atom molecular dynamics (MD) trajectories. For α-synuclein, a 30 µs MD simulation using an advanced force field ^6^ failed to reproduce the transient N-terminal α-helix observed experimentally, whereas AI-IDP captured this feature directly from sequence. Only when MD was combined with a hierarchical chain-growth strategy did transient helicity emerge, at substantially higher computational cost (Extended Data Fig. 15). These comparisons highlight that accurate description of IDP ensembles can be achieved without long-timescale simulations, demonstrating that AI-IDP encodes key physical determinants of disorder within a data-driven yet efficient framework.

We note that deviations between predicted and experimental secondary structure propensities can arise from environmental modulation of local structure. For instance, Cu(I) binding stabilizes α-helical conformations at the N-terminus of α-synuclein,^20^ increasing experimental SSP values beyond those predicted by AI-IDP (Extended Data Fig. 10). Such cases underscore that AI-IDP captures intrinsic, sequence-encoded propensities, while environmental factors such as metal or lipid binding can further stabilize transient structure in specific cellular contexts.

Together, these results demonstrate that AI-IDP quantitatively recovers the sequence-dependent transient structure of IDPs. The method therefore establishes a direct link between amino-acid sequence and local conformational bias - a key determinant of IDP function and molecular recognition.

### Medium-range contacts and chain organization

We next asked whether AI-IDP captures transient medium-range contacts that influence the overall organization of intrinsically disordered proteins. Such contacts, typically 10-40 residues apart, are critical for modulating chain compaction and molecular recognition and are experimentally probed by paramagnetic relaxation enhancement (PRE) measurements ^21^. For α-synuclein, PRE profiles measured with spin labels at either Cys18 (N-terminus) or Cys140 (C-terminus) closely matched those back-calculated from AI-IDP ensembles (Fig. 1B (viii)). The predicted and experimental profiles showed close agreement, including the characteristic attenuation shoulder around residues 30-70 observed for the Cys18-labelled sample. Inspection of the AI-IDP ensemble revealed that this feature arises from transient contacts between the N-terminal and central regions, mediated by a flexible Ala-Gly linker. In contrast, both experiment and AI-IDP predictions indicated weaker medium-range interactions when the label was placed at the C-terminus (Fig 1B (ix)), consistent with the known chain expansion in this region ^22^.

To test a distinct system, we analyzed the repeat domain of Tau (fragment K18), for which experimental PREs had revealed transient compaction involving the hydrophobic hexapeptides ^275^VQIINK^280^ and ^306^VQIVYK^311^. PRE profiles back-calculated from Tau’s AI-IDP ensemble reproduced these features, showing enhanced broadening at the spin-label sites 260 and 291 and identifying short-lived contacts between the hexapeptides (Fig. 1B (x)). Structural inspection linked these interactions to local turns in the preceding PGGG motifs (^271^PGGG^274^ and ^301^PGGG^304^), which bend the chain and facilitate encounter between repeats. Replacing the glycines with prolines in silico disrupted these turns and increased the average end-to-end distance of the segment (residues 291-311) by approximately 4 Å, confirming the sequence dependence of Tau chain compaction. These results demonstrate that AI-IDP not only reproduces transient local structure but also captures sequence-encoded medium-range contacts that shape IDP architecture. The ability to detect such subtle interactions - without explicit force-field parameterization - suggests that AI-IDP effectively integrates local conformational bias with medium-range organization. This is also supported by the finding that PRE values predicted from AI-IDP ensembles demonstrate high correlations with experiment and low RMSD values (Extended Data Fig. 16; 14 IDP/IDRs in Extended Data Table 4). AI-IDP PRE RMSDs are only slightly higher than those of idpGAN ensembles, which however display low agreement with experimentally observed global chain dimensions (Extended Data Fig. 19)

### Local structure determines compaction

Having shown that AI-IDP captures transient local and medium-range structure, we next asked whether such sequence-encoded local biases are sufficient to explain global dimensions of disordered proteins. To this end, we generated AI-IDP ensembles for 137 IDPs with experimentally determined radius of gyration (*R*_g_) values from small-angle X-ray scattering (SAXS) (Fig. 1C) ^23^.

Across this diverse dataset, AI-IDP accurately reproduced global chain dimensions, with an overall correlation coefficient of 0.95 and a root-mean-square deviation of 3.6 Å between predicted and experimental *R*_g_ values (Fig. 1C (xi, xiii)). These ensembles, generated without explicitly incorporating long-range chain interactions (e. g. electrostatic or hydrophobic/aromatic), thus capture the effective compaction of IDPs directly from sequence information. A comparison with CALVADOS ^14^, a coarse-grained method that explicitly models long-range interactions, yielded a similar correlation (r = 0.96) but lacked the experimentally observed transient helices and medium-range contacts present in AI-IDP ensembles (Extended Data Figs. 11, 17, 19). On the other hand, spuriously high SSPs in IDPFold predictions of several IDPs, reflecting inaccuracies in transient structure predictions, likely also result in spurious intrachain contacts (PRE plots C-F in Extended Data Fig. 18) and less optimal agreement between predicted and experimental *R*_g_ values (Extended Data Fig. 19).

To explore how local conformational preferences contribute to overall dimensions, we analyzed the population of residues adopting polyproline-II conformations within each AI-IDP ensemble. Polyproline-II helices, known to extend peptide backbones, were strongly associated with global chain expansion: ensembles with higher polyproline-II content exhibited larger *R*_g_ values, with correlation coefficients of 0.42 for predicted and 0.39 for SAXS-derived compaction indices (Fig. 3). IDPs containing more than 10% polyproline-II residues showed statistically significant increases in chain expansion relative to those with lower polyproline-II content.

**Figure 3.**
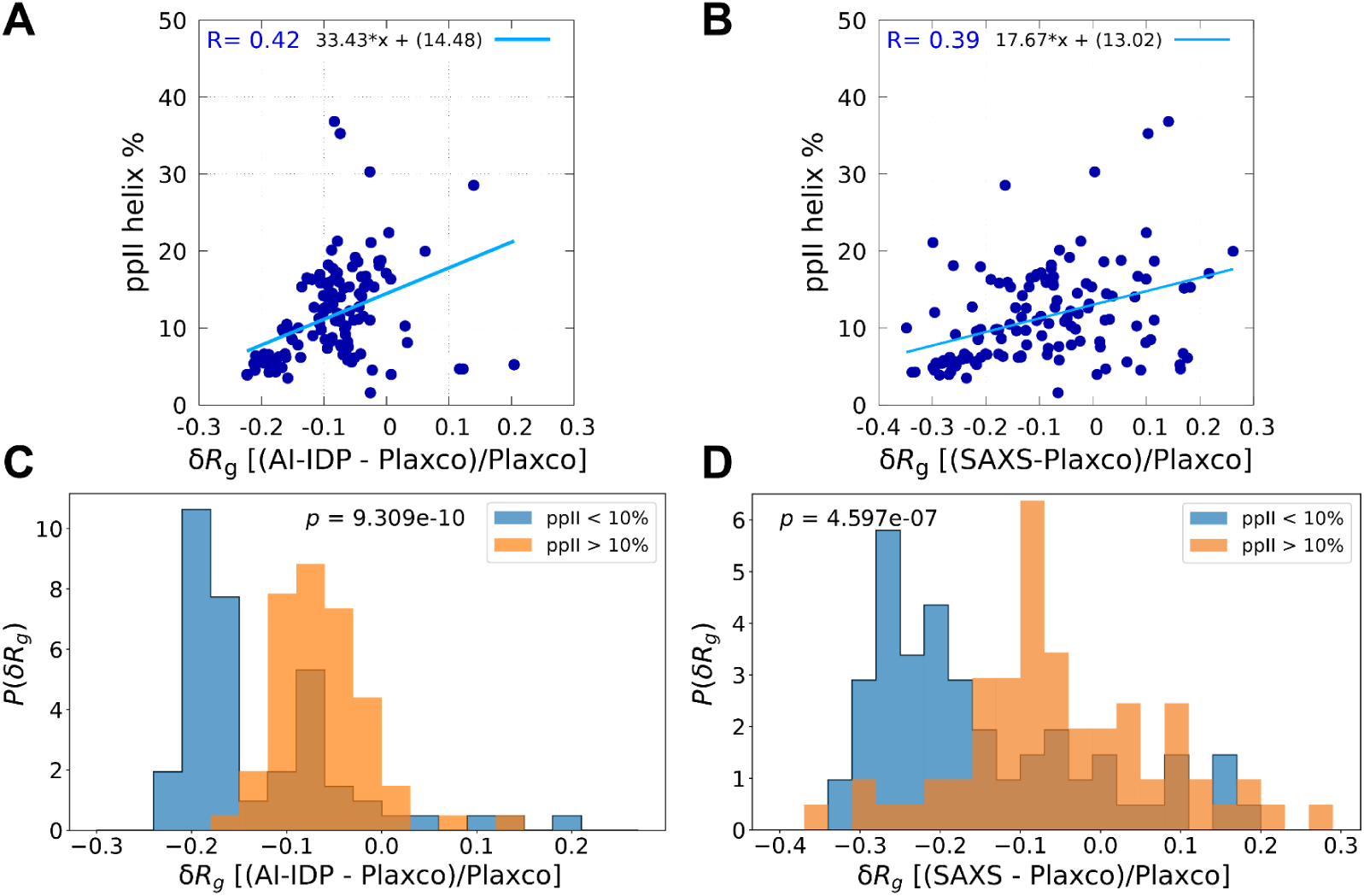
Transient polyproline-II structure drives IDP expansion. **A, B.** Correlation between the predicted average percentage of residues in polyproline-II conformation in AI-IDP ensembles and chain expansion of 137 IDPs, calculated as a relative difference (δ*R*_g_) between predicted (A) or experimental, SAXS-derived *R*_g_ (B) and *R*_g_ of a random-coil chain (Plaxco value). **C, D.** Distributions of AI-IDP-derived (C) and SAXS-based (D) chain expansions (δ*R*_g_) for proteins with polyproline-II conformational content below and above 10% among 137 disordered proteins. *p*-value (obtained from t-distribution using SciPy v1.16) corresponds to one-sided Brunner-Munzel test comparing δ*R*_g_ from both groups of proteins.

Both AI-IDP and CALVADOS-generated ensembles demonstrate lower RMSDs between calculated and experimental *R*_g_ values than Flexible-Meccano which incorporates structural propensities only at a residue level (Extended Data Fig. 19). These findings indicate that the overall dimensions of IDPs can emerge not only from long-range chain interactions (present in CALVADOS ensembles) but also from the interplay of sequence-encoded local structure and transient medium-range contacts (present in AI-IDP ensembles). In particular, the prevalence of extended polyproline-II helices and flexible linkers provides a physical basis for the chain expansion observed in disordered proteins. By linking sequence composition to global conformational behaviour, AI-IDP establishes a framework linking local and global scales of IDP organization.

### Conformational ensembles of giant IDPs

A major challenge in studying intrinsically disordered proteins is that many biologically central examples are thousands of residues long and remain inaccessible to high-resolution experimental analysis. We therefore applied AI-IDP to several “giant” IDPs to test whether the approach can resolve their sequence-dependent structural organization.

Titin, the principal elastic element of human muscle, contains a 2152-residue disordered region rich in proline, glutamate, valine, and lysine. In its AI-IDP ensemble, more than half of the residues adopt polyproline-II conformations (Fig. 4A), providing a plausible molecular origin for Titin’s extensibility ^24^. Two short helical segments were also identified, both conserved among mammals and located in sequence regions implicated in protein-protein interaction (Extended Data Table 5). These elements may locally tune the mechanical properties of Titin while providing potential recognition motifs.

**Figure 4.**
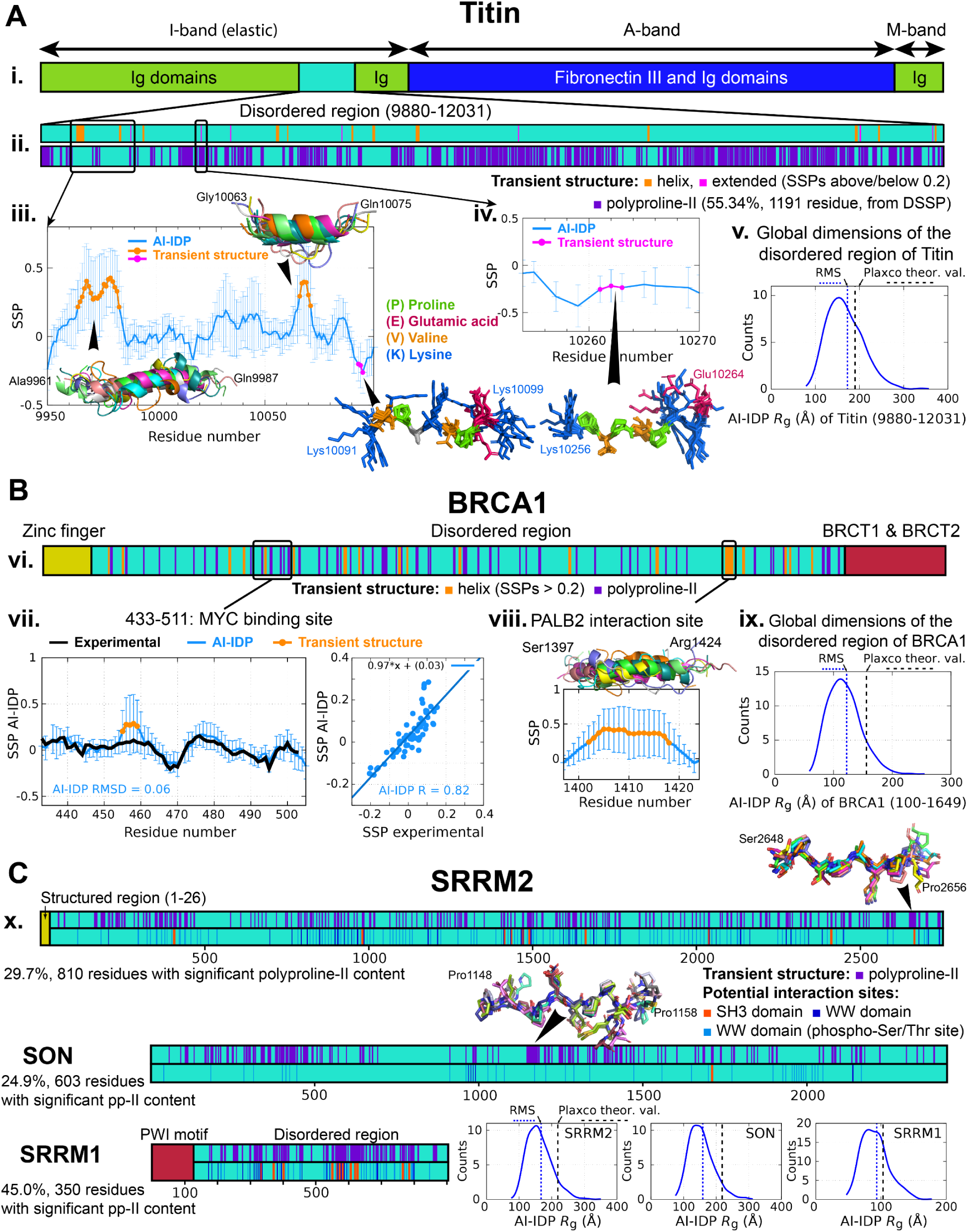
Decoding conformational heterogeneity in giant intrinsically disordered proteins. **A.** Titin. (i) Domain organization highlighting the 2,152-residue disordered region (residues 9,880–12,031). (ii) Sequence-resolved map of transient secondary structure in the AI-IDP ensemble: α-helical (orange), extended (pink), and polyproline-II (ppII, purple) propensities. (iii-iv) Enlarged views of representative disordered segments showing transient helices and extended regions identified from AI-IDP-derived SSP scores. (v) Distribution of radii of gyration (*R*_g_) across Titin’s AI-IDP ensemble, compared with the ensemble-averaged (RMS) and theoretical random-coil *R*_g_ value. **B,** BRCA1. (vi) Sequence-specific map of transient structure from the AI-IDP ensemble, with known interaction sites for Myc and PALB2 indicated. (vii) Residue-specific SSP scores for the Myc-binding region derived from experimental NMR data (black) and the AI-IDP ensemble (blue); correlation plot at right. (viii) Representative AI-IDP conformers illustrating the transient α-helix that mediates the BRCA1-PALB2 interaction. (ix) *R*_g_ distribution of the BRCA1 ensemble with ensemble-averaged and theoretical random-coil values. **C,** RS-rich nuclear speckle proteins. Sequence-resolved maps of polyproline-II structure in the AI-IDP ensembles of SRRM2, SRRM1, and SON. Predicted SH3- and WW-domain interaction motifs are indicated below, with phosphorylation-regulated WW-binding sites in light blue. Lower right, distributions of *R*_g_ values for SRRM2, SRRM1, and SON compared with theoretical expectations. Together, the panels demonstrate that AI-IDP scales to the largest known disordered proteins, capturing conserved α-helical and polyproline-II motifs that underlie elasticity, molecular recognition, and phase-separation behavior in giant IDPs.

We next examined BRCA1, a tumor suppressor that orchestrates DNA damage repair ^25^. BRCA1 illustrates how AI-IDP can rationalize functional motifs embedded within long disordered regions. Predicted secondary-structure propensities within BRCA1’s AI-IDP ensemble agreed with experimentally derived SSP profiles, particularly at residues corresponding to the Myc-binding region (Fig. 4B). In addition, AI-IDP correctly identified the transient α-helical segment that mediates BRCA1/PALB2 complex formation (Fig. 4B), whose stability is essential for homologous recombination repair ^26^.

Finally, we analyzed the arginine/serine-rich (RS)-rich nuclear proteins SRRM2 and SON, which form the architectural core of nuclear speckles ^27^. Both proteins exhibited distinctive disorder signatures: their AI-IDP ensembles contained extensive polyproline-II-rich regions accounting for approximately 25-30% of residues, consistent with their proline-enriched composition (Fig. 4C). The shorter homolog SRRM1 displayed an even higher polyproline-II content (∼45%), suggesting that polyproline-II helices contribute to the weak, multivalent interactions that promote nuclear-speckle condensation and spliceosome assembly ^28,29^.

Together, these results demonstrate that AI-IDP scales to the largest known disordered proteins, generating interpretable, sequence-resolved ensemble models far beyond the reach of conventional experimental or simulation approaches. The predicted enrichment of polyproline-II conformations across giant IDPs provides a physical basis for their elasticity, solubility, and phase-separation behavior, while conserved α-helical motifs highlight potential interaction hotspots.

### Sensitivity to point mutations

Because intrinsic disorder often mediates regulation through sequence-encoded conformational biases, even single amino-acid changes can remodel ensemble structure and function. To test whether AI-IDP captures such effects, we analysed disease- and modification-linked variants in two representative systems: the RNA-binding protein TDP-43 and the transcription factor c-Myc.

The C-terminal domain of TDP-43 (residues 267–414) undergoes liquid–liquid phase separation, and mutations that alter its transient structure modulate aggregation propensity ^30^. Two amyotrophic lateral sclerosis (ALS)-associated variants, A321G and A326P, disrupt the short α-helix (residues 310–350) that stabilizes intermolecular contacts in the wild-type protein. AI-IDP ensembles reproduced this behaviour: secondary-structure propensities correlated strongly with NMR data (r = 0.85, RMSD = 0.09), and the predicted helical population within the 310-350 segment decreased from 14.8 % in wild-type to 12.1 % (A321G) and 5.7 % (A326P) (Fig. 5A). These reductions mirror the experimentally observed loss of helical content and diminished phase-separation capacity.

**Figure 5.**
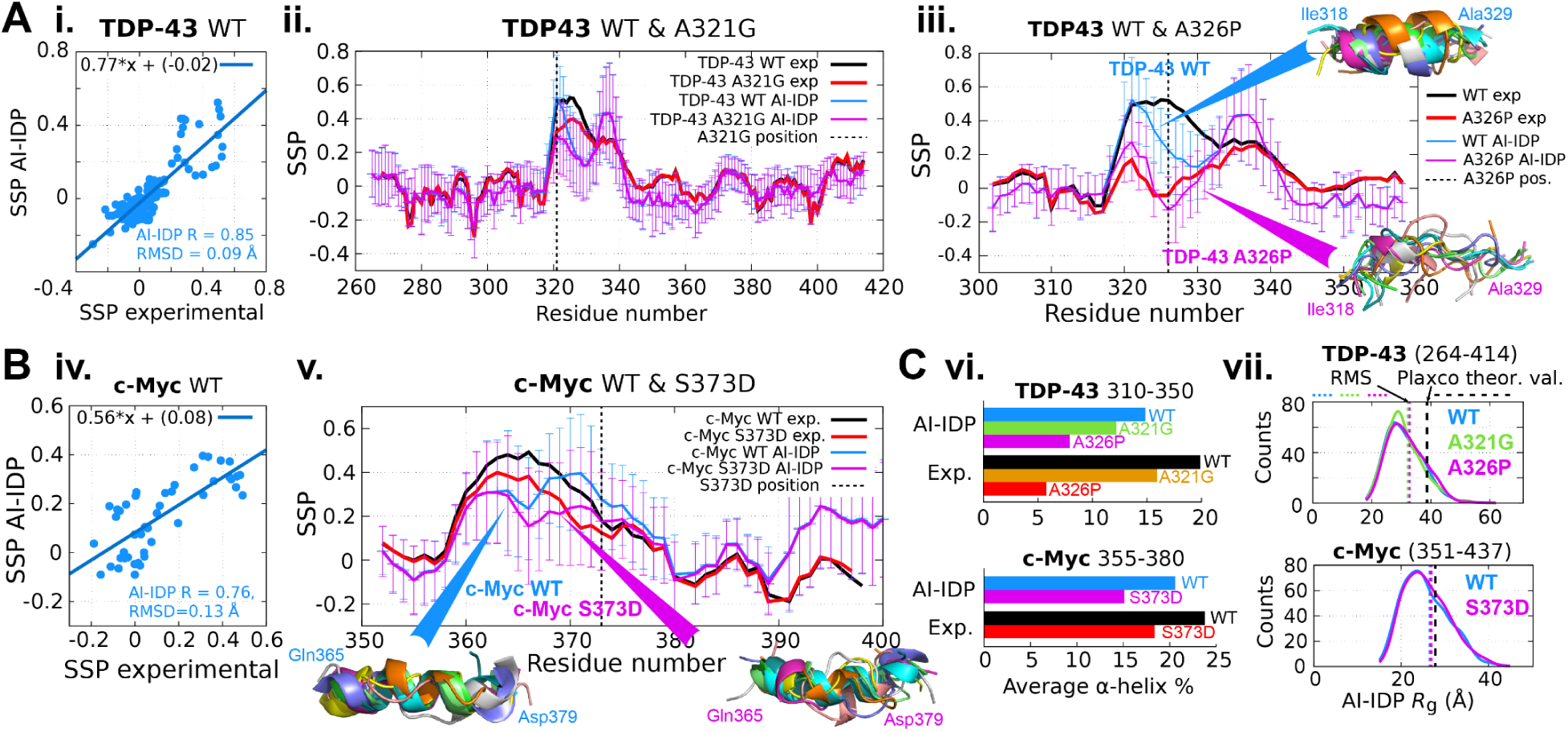
AI-IDP ensembles reproduce the structural impact of mutations in intrinsically disordered proteins. **A. TDP-43.** (i) Correlation between residue-specific SSP scores derived from the AI-IDP ensemble of wild-type TDP-43 (residues 267–414) and experimental NMR SSP scores. (ii, iii) Comparison of AI-IDP-predicted and experimental SSP profiles for the C-terminal domain of wild-type (blue and black lines, respectively), A321G and A326P variants (red and pink lines), showing loss of transient helicity in the 310–350 region. **B. c-Myc.** (iv) Correlation between experimental and AI-IDP-predicted SSP scores for wild-type c-Myc. (v) Residue-specific SSP profiles for wild-type and phosphomimetic S373D c-Myc, showing local disruption of the transient α-helix (residues 365–379); representative AI-IDP conformers are shown below. **C.** (vi) Fractional α-helical population in a segment of TDP-43 and c-Myc from experimental data and AI-IDP ensembles; both TDP-43 variants and c-Myc variant display reduced helicity relative to wild type. (vii) Distribution of radii of gyration (*R*_g_) in wild-type and mutant ensembles, with ensemble-averaged and theoretical random-coil values indicated. Together, these data show that AI-IDP quantitatively captures the effects of single-residue substitutions and post-translational mimetics on the conformational ensembles of disordered proteins, linking sequence perturbations to measurable structural changes.

We next examined c-Myc, whose disordered basic helix-loop-helix region transiently forms an α-helix that binds the partner Max. Phosphorylation of Ser373 disrupts this motif and weakens Max binding ^31^; the phosphomimetic S373D variant reproduces this effect in vitro. AI-IDP reproduced both the wild-type transient helix and its local destabilization upon mutation (Fig. 5B), in agreement with NMR-derived SSP profiles. The predicted reduction in helical propensity explains the experimentally observed decrease in binding affinity.

By contrast, other ensemble generators lacked this sensitivity: CALVADOS and IDPConformerGenerator produced static or over-helical ensembles and failed to reproduce mutation-induced structural changes (Fig. 5B and Extended Data Fig. 20). The ability of AI-IDP to detect such subtle shifts arises from its hybrid design, sequence-conditioned fragment prediction combined with flexible physical assembly, which preserves the mutational context of local dihedral distributions.

These findings demonstrate that AI-IDP quantitatively resolves the impact of point mutations and post-translational modifications on disordered-protein ensembles, directly linking sequence perturbations to structural outcomes. By capturing how minimal changes in amino-acid composition reshape transient structure, AI-IDP provides a mechanistic framework for interpreting variant effects, understanding disease-associated conformational shifts, and guiding the rational design of modulators targeting dynamic disordered regions.

### Evolution of transient structure in disordered proteomes

Structural disorder is pervasive across life, yet its evolutionary patterns remain poorly understood because conformational ensembles have been determined for only a few short IDPs. To provide a quantitative view of how transient structure has diversified across evolution, we generated AI-IDP ensembles for all experimentally confirmed disordered regions in the DisProt database ^32^, encompassing more than 3,000 proteins and over 290,000 residues from human and non-human organisms (Fig. 6A; Extended Data Fig. 21). Analysis of these ensembles revealed consistent sequence-structure trends: while α-helicity showed organism- and function-specific variation, polyproline-II content increased systematically from prokaryotes to higher eukaryotes. The prevalence of transient secondary structure varied widely across proteins, with some disordered regions exceeding 80 % α-helical or 60 % polyproline-II propensity (Fig. 6B-C).

**Figure 6.**
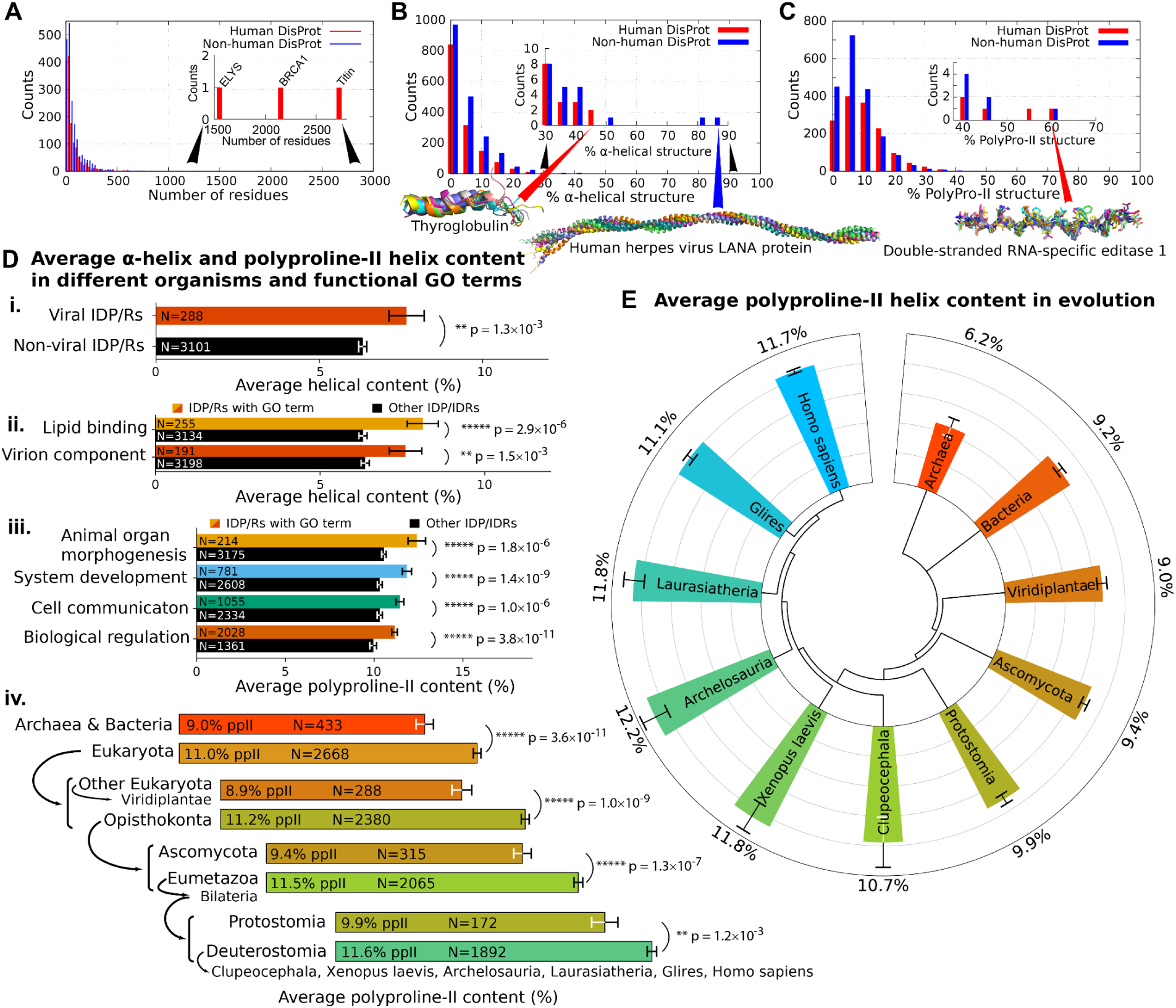
Evolution of transient structure in intrinsically disordered proteomes. **A.** Distribution of sequence lengths for intrinsically disordered proteins (IDPs) annotated in the DisProt database. **B, C,** Proteome-wide abundance of transient α-helical and polyproline-II conformations predicted by AI-IDP across all DisProt entries; representative ensembles illustrating highly α-helical and polyproline-II-rich regions are shown. **D,** Functional and taxonomic trends in transient structure. (i,ii) Viral IDPs show higher α-helical content than those from other organisms, reflecting their pre-formed recognition motifs. (ii) IDPs associated with lipid binding exhibit elevated α-helical propensity, consistent with membrane association. (iii) IDPs involved in organism development, cell communication and regulation are enriched in polyproline-II conformations characteristic of proline-rich, multivalent interaction motifs. (iv) Average polyproline-II content increases significantly at multiple evolutionary branch points (p < 0.05, Brunner-Munzel test), indicating progressive expansion of polyproline-II bias from prokaryotes to higher eukaryotes. **E,** Phylogenetic tree of taxa represented in DisProt, annotated with the mean polyproline-II content of their AI-IDP ensembles, illustrating evolutionary tuning of conformational bias. Together, these analyses reveal that transient α-helices and polyproline-II segments are universal but differentially enriched motifs whose balance has diversified through evolution to support increasing complexity in signaling and regulation.

We next examined how these conformational tendencies relate to function and taxonomy. Viral IDPs showed a pronounced enrichment in transient α-helices (Fig. 6D (i), Extended Data Table 6), consistent with their need to engage diverse host factors using short, preformed motifs. Among functional gene ontology categories, IDPs linked to lipid binding also exhibited higher α-helical propensities (Fig. 6D (ii), Extended Data Table 6), suggesting that partial pre-structuring facilitates membrane association. By contrast, IDPs linked to functions important in organism development and regulation displayed elevated polyproline II content (Fig. 6D (iii), Extended Data Tables 6, 8), reflecting their proline-rich, low-complexity composition and propensity for dynamic, multivalent interactions.

Across taxa, AI-IDP ensembles revealed a progressive increase in polyproline II content from prokaryotes to higher eukaryotes (Fig. 6D (iv); 4E; Extended Data Table 7). This shift parallels the expansion of signaling and regulatory networks in multicellular organisms and is consistent with the evolution of sequence-encoded conformational bias to support more complex modes of transient recognition. The RS-rich scaffold proteins of nuclear speckles, SRRM2 and SON, exemplify this trend: both lengthened and accumulated proline-rich segments during evolution ^27^, increasing their predicted polyproline II helicity and likely enhancing their capacity for phase-separated assembly.

Together, these analyses reveal that the conformational properties of IDPs are not uniform but evolutionarily tuned. Transient α-helices and polyproline II segments emerge as widespread structural motifs whose prevalence and balance reflect adaptive trade-offs between recognition specificity and dynamic flexibility. By extending ensemble prediction to the proteome scale, AI-IDP provides a framework to trace how sequence-encoded disorder has diversified across evolution and how this diversification underpins the structural logic of cellular regulation.

### Conclusions

AI-IDP bridges the gap between static protein structure prediction and the dynamic reality of intrinsically disordered proteins by transforming sequence information into realistic conformational ensembles. Integrating deep-learning-based fragment prediction with flexible physical assembly, it achieves ensemble-level accuracy from local structure to global dimensions and across entire proteomes by providing accurate local fragment information. The resulting landscapes reveal that transient α-helices and polyproline-II structures are pervasive, evolutionarily tuned features of disorder that link sequence to function and regulation. While environmental context, the presence of folded domains, multichain coupling and ligand binding remain to be incorporated, the framework provides a practical solution for decoding conformational heterogeneity directly from sequence. By uniting artificial intelligence with physical insight, AI-IDP establishes a foundation for systematic exploration of protein dynamics and for the rational design of modulators targeting the disordered proteome.

## Acknowledgements

We thank Alain Ibáñez de Opakua for designing the initial version of the fragment assembly algorithm, Lukas S. Stelzl and G. Hummer for making available the MDS HCG ensemble of α-synuclein, Paul Robustelli and D.E. Shaw for the MD-trajectory of full-length α-synuclein, and Marco C. Miotto and Claudio O. Fernandez for the chemical shifts of Cu(I)-bound α-synuclein. A.A. and M.Z. are grateful for computational support and infrastructure provided by the Max Planck Computing and Data Facility (MPCDF). M.Z. was supported by the European Research Council (ERC) under the EU Horizon Europe research and innovation programme (project No. 101141570).

## Author contribution

A.A. built the AI-IDP algorithm for the generation of conformational ensembles, generated AI-IDP ensembles, and analyzed their structural features. M.Z. designed the project. A.A. and M.Z. wrote the manuscript.

## Ethics declarations

A. Abyzov and M. Zweckstetter are employees of brainQr Therapeutics GmbH.

## Methods

### Structural ensemble calculations

The process of determining ensembles of structures for IDPs/disordered regions is schematically illustrated in Extended Data Fig. 1. The process starts with the generation of AlphaFold2 structures for all possible fragments of fixed length. Subsequently, full protein chains are assembled from these fragments using an in-house Pymol script.

For each IDP sequence, 10-residue fragments were generated covering all sequence with one-residue shift between fragments. For example, for a 100-residue IDP, 91 fragments were generated: 1-10, 2-11… 91- 100. Five structures (ranked from 1 to 5) were predicted for each 10-residue fragment using the Colab implementation of AlphaFold (ColabFold ^1^, AlphaFold2 w/MMseqs2, BATCH mode) on a Google v2 TPU or Google v6 TPU VM. The ColabFold parameters were msa_mode=MMseqs2 (UniRef+Environmental), num_models=5, num_recycles=3, stop_at_score=100, use_amber=yes, use_templates=no. We used structures produced after the Amber relaxation step. For an IDP with N residues, we generated 5*(N-9) fragment structures.

From these AlphaFold-predicted structures of 10-residue fragments, full-length structures of the AI-IDP ensembles were generated with an in-house Python script using open-source Pymol (v3.0.0) (Extended Data Fig. 1, i.). Each successive fragment is shifted by 8 residues in sequence compared to the previous fragment, so that two residues between each fragment pair would overlap. The two first fragments (residues 1..10 and 9..18) are aligned by aligning atoms C_α_, C’ of residue 9 and atom N of residue 10 (representing the peptide bond between these residues) of the first fragment with corresponding atoms in the second fragment (Extended Data Fig. 1, ii.). Residue 10 of the first fragment and residue 9 of the second fragment are dropped. After that, we introduced flexibility in the newly formed link between fragments randomly changing the dihedral ϕ and ψ angles. If Pymol detected a helical structure in either residue 9 or 10, dihedral angles were not modified to preserve the helix. However, in case of significant clashes between fragments (any 2 heavy atoms with distance < 1.18 Å) we modified dihedral angles anyway to resolve these clashes.

Dihedral angles were modified according to the following procedure. We first set ω (peptide bond) angles between residues 7-11 to 180°. Then we sampled backbone dihedral ϕ and ψ angles in residues 8-10 from a library of amino acid-specific angles obtained from unstructured regions of high-resolution X-ray structures ^2^. After sampling each pair of angles, we checked the C_α_^i^,C_α_^i+1^ – C_α_^i+4^, C_α_^i+5^ angle formed between vectors connecting C_α_ atoms of residues 7-8 and 10-11. We continued sampling until this angle was below an empirically-determined threshold of 40° (Extended Data Table 1) or we had sampled 100 ϕ/ψ pairs, whichever came first (Extended Data Fig. 1, iii.). If significant clashes were found in the whole chain (any two heavy atoms with distance < 1.18 Å), we repeated the sampling procedure from the beginning (iv).

Subsequent fragments were added to the nascent chain following the same steps as for connecting the first two fragments. To increase conformational sampling, the starting point for the first 10-residue fragment was chosen from one of 8 first residues in the sequence (Extended Data Fig. 1, ii.), and the structure of each fragment was randomly chosen from one of five alternatively predicted structures. If the starting point of the chain was not the first residue in the IDP sequence, we added a fragment covering the first 10 residues in the sequence to fill in the N-terminal gap before joining first two fragments as discussed above. Finally, we added a fragment covering the last 10 residues in the sequence to avoid a potential C-terminal gap in the formed chain. The formed chain had no significant clashes because for each fragment joining operation, we sampled ϕ/ψ angle pairs in a way that prevented these clashes. We removed hydrogen atoms from the generated structures as their positions are not precise in AlphaFold-predicted fragment structures, because AlphaFold was trained on PDB structures mostly coming from X-ray crystallography or cryo-EM experiments where protons are often incorporated by computational modelling.

For each IDP, we generated 1000 structures, in line with recent publications ^3,4^. Our tests showed that around 100-200 AI-IDP structures (for SSPs and ^3^J(H_N_H_α_) couplings) and around 500-1000 structures (for PREs and *R*_g_) are required for the convergence of ensemble averages of calculated parameters (Extended data Fig. 3).

We generated structures using 6-, 8-, and 12- residue fragments in an equivalent way. We calculated C_α_ and C_β_ chemical shifts and secondary structure propensities (SSPs) from ensembles of structures generated with different fragment sizes, and compared these SSP scores with experimentally-derived ones. By minimizing the root mean square deviation (RMSD) and maximizing Pearson’s r correlation coefficient between experimental and calculated SSPs, we determined the optimal fragment length as 10 residues (Extended Data Fig. 2). In the same way, we determined that using the most powerful sequence alignment option proposed by ColabFold (MMseqs2 UniRef+Environmental) resulted in a better prediction of IDP transient structure, as illustrated by residue-specific SSP scores, RMSD and Pearson’s r values (Extended Data Fig. 4).

We also evaluated the effect of random choice of one of five structures generated for each fragment or the randomly picked order of residue-specific ϕ/ψ angles used for fragment link flexibilization. These did not result in any visible differences in derived SSP profiles or *R*_g_ values (Extended Data Fig. 5).

Importantly, only the fragment generation was performed by a deep-learning model AlphaFold2 which already received extensive training and validation. The subsequent fragment assembly procedure did not involve any AI models that would require training or validation, therefore eliminating risks coming from deficiencies in these procedures.

### Ensemble analysis

For each of the 1000 generated structures, C_α_ and C_β_ chemical shifts were predicted using UCBShift ^5^. We calculated residue-specific SSP scores ^6^ from predicted as well as from experimental C_α_, C_β_ chemical shifts. Predicted SSP scores were averaged over an ensemble of structures. For comparison purposes, we also predicted chemical shifts using SPARTA+ ^7^ and SHIFTX2 (1.10A) ^8^ and calculated SSPs from them (Extended Data Fig. 6). We compared SSPs from experimentally obtained and ensemble-predicted C_α_, C_β_ chemical shifts and calculated root mean square deviation (RMSD) values and Pearson’s r correlation coefficients (Extended Data Fig. 6). We concluded that UCBShift was performing better than both SPARTA+ and SHIFTX2, as it both better reproduced regions of elevated experimental SSPs (e.g. in cMyc residues 355-380), and its use resulted in overall smaller RMSD values and higher correlation coefficients between experimental and ensemble-calculated SSPs.

Even though SSPs are indirect indicators of secondary structure, we can directly compare them with SSPs derived from experimental chemical shifts and evaluate the accuracy of structural feature prediction in IDPs. We calculated a sequence average of positive-only SSPs (x > 0? yes: x; no: 0), representing the average amount of transient α-helical structure ^6^ in an IDP, and correlated it with an average (over sequence and ensemble) amount of transient α-helical structures described by the DSSP algorithm ^9–11^ in the generated ensemble of structures. For SSP scores calculated from UCBShift-predicted chemical shifts, we obtained a Pearson correlation factor of 0.9 and a close to 1:1 linear relationship, supporting the use of SSP scores as secondary structure indicators. For chemical shifts predicted by SPARTA+ the correlation was significantly lower (R = 0.5) and SSP-determined helical propensities were lower than those determined by DSSP (Extended Data Fig. 7).

Whenever three or more consecutive residues in an IDP sequence had absolute SSP scores above 0.2, we defined these residues as possessing significant transient structure (α-helical for positive SSPs and β-sheet/extended for negative SSPs) ^6^.

To calculate ^3^J-H_N_H_α_ couplings, we added protons to the ensemble of 1000 generated structures using fixPDB.tcl tool from NMRPipe package ^12^, and subsequently calculated θ = ∠(H_N_, N, C_α_, H_α_) dihedral angle in Pymol and coupling value ^3^*J* H_N_H_α_ = 6.51 cos^2^ θ − 1.76 cos θ + 1.6.

To calculate paramagnetic relaxation enhancement (PRE) line broadening for predicted ensembles with added protons, we employed a modified version of the PREdict Python tool from DEERpredict ^13^. As the original version predicted overly extensive line broadening around spin label positions, we replaced Boltzmann weights, associated with the probability of each spin label rotamer conformation in PREdict, by a simpler 1/0 function to avoid steric hindrance. This function was equal to 1 if the minimal distance between the center of line connecting N and O atoms of spin label and the position of backbone amide atoms was equal to or above 5.0 Å, otherwise 0. We used the “MTSSL 175K CASD” rotamer library. Other parameters were *τ*_c_ = 5 ns, *τ*_i_ = 200 ps, total INEPT-delays in the pulse sequence were 10 ms, *R*_2,dia_ = 12.566 s^-1^. Temperature was 288 K and proton Larmor frequency was 600 MHz for α-synuclein, 278 K and 800 MHz for Tau K18 and 278 K and 900 MHz for full-length human tau (htau40).

Experimental NMR chemical shifts were obtained either from the BMRB database for 4E-BP2 (code 19114), ACTR (code 15397), BRCA1 219-504 (code 50231), cMyc 351-437 WT (code 27414), cMyc 351-437 S373D (code 27418), p53 (code 51984), TDP-43 268-414 WT (code 26823), TDP-43 268-414 A321G (code 26826), TDP-43 268-414 A326P (code 26828) or from published papers for α-synuclein ^14^ and from the work of our group on tau protein ^15^. Experimental ^3^J-H_N_H_α_ couplings were obtained from the work of our group on α-synuclein ^14^. Experimental PRE profiles were obtained from published papers on α-synuclein ^16–18^, Tau K18 ^19^ and full-length tau ^20^.

### Comparison with different IDP ensemble generation methods

To compare the performance of AI-IDP with other recently published IDP conformational ensemble generation methods, we generated ensembles using CALVADOS ^3^, IDPConformerGenerator ^4^, idpGAN ^21^, Flexible-Meccano ^22^, IDPFold^23^, IDPFold2^24^, IDP-o^25^ and STARLING^26^ for 40 different IDPs (Extended Data Table 2-2) to compare SSPs, and using CALVADOS, idpGAN, IDPFold and STARLING for 14 different IDPs (Extended Data Table 2-3) to compare PREs. Each ensemble had 1000 structures, except for IDPFold, where 576 structures were generated for SSP and *R*_g_ comparisons, and twice as much for PRE comparisons. CALVADOS ensembles (CALVADOS2 version) were generated using the IDRLab Colab notebook and side chains were added using the PULCHRA software ^27^ directly in the notebook workflow. Temperature and ionic strength were matched with the experimental conditions of the NMR data. IDPConformerGenerator ensembles were generated using sampling without biasing for secondary structure (--dany --dloop-off) and with MCSCE algorithm ^28^ for side chain building. For htau40, we used IDPConformerGenerator ensemble of 1000 structures from the Protein Ensemble Database (PED, proteinensemble.org). idpGAN coarse-grained ensembles were generated using idpgan_ped python package^29^ notebook and all-atom structures were generated using the cg2all software ^30^, and relaxed using amber force fields as in ColabFold ^1^. Same reconstruction and relaxation was applied to STARLING and IDPFold2 ensembles. For IDPFold ensembles, fine-tuned model was used, side chains were reconstructed using the pdbfixer^31^ software and subsequently relaxed with amber. For STARLING ensembles, ionic strength was matched with the experimental conditions of the NMR data. Flexible-Meccano (latest version from 16 October 2012) ensembles were generated with no secondary structure bias and side chains were added using the FASPR algorithm ^32^. For Rg comparisons, IDPFold and Flexible-Meccano ensembles were calculated as previously described, and values for CALVADOS and idpGAN were obtained from ref.^33^ Cα/Cβ chemical shifts, SSP scores and where applicable PRE profiles were calculated in the same way as for AI-IDP ensembles.

### Runtime

The ColabFold runtime was ∼1 minute per fragment using Google Colab v2-8 TPU, from which 15 seconds was the actual calculation time and remaining 45 seconds was the average waiting time for Multiple Sequence Alignment (MSA) requests from the public MSA server. The Python/Pymol script chain generation runtime was on average 0.06*x^2^ seconds for 1000 structures of an IDP with x residues on Intel® Core™ i7-13700H laptop. Typical running times for AI-IDP and other IDP ensemble generation methods are presented in Extended Data Table 7. UCBShift runtime for 1000 structures of the 140-residue α-synuclein, when only C_α_ and C_β_ chemical shifts were calculated, was ∼30 minutes on the same Intel laptop with 4 parallel instances of UCBShift running. For 441-residue htau40, it was ∼85 minutes. We also performed chemical shift predictions on Google Colab (TPU v2-8 runtime) with 2x 24-core Intel® Xeon® CPUs @ 2.00GHz. UCBShift runtime (50 parallel instances) for 1000 structures of α-synuclein with only C_α_, C_β_ chemical shifts calculated was 7 minutes, whereas for htau40 it was 22 minutes.

### Giant IDP/IDRs analysis

To find regions in giant IDPs (Figure 2) with predicted transient secondary structure, we determined where in the sequence three or more consecutive residues had SSP scores (calculated as above) of more than 0.2 – defining these residues as part of transient α-helical structure, and less than -0.2 – defining these residues as part of transient β-sheet/extended structure. For polyproline-II helical content, we used the DSSP algorithm ^9–11^ (version 4) to assign secondary structure propensity for each residue in each member of the generated ensemble. When more than 20% of structures in the ensemble were assigned polyproline-II helical conformation at a particular residue, that residue was defined as part of a transient polyproline-II helical region. We subsequently calculated percentages of such residues in the IDP/disordered region sequence.

Potential SH3 domain interaction sites were identified on the basis of following sequence motifs: [R/K]XXPXXP or PXXPX[R/K]. Potential WW domain interaction sites were identified based on following sequence motifs: PPXY, LPXY, PPR, PGM or PR.

### Selection of intrinsically disordered proteins and regions

For the proteome analysis, we downloaded all entries from the DisProt release 2024_06 (JSON format). We treated each entry as IDP/disordered region that was assigned the term “disorder” in the “structural state” category of the DisProt consensus track, merging together all regions that had no gap between them. We retained only disordered regions longer than 15 residues, which resulted in 3389 sequences from 2604 different proteins containing in total 292910 residues. Each disordered region in the resulting database was assigned a name on the bases of its UniProt ID and first and last residues, separated by dots (e.g. P04637.1.93).

### Proteome analysis

In the distribution plots of the number of IDPs/disordered regions with a given amount of α-helical or polyproline-II structure (Figure 4 B-C), the latter was calculated as an average percentage of residues in the generated conformational ensemble assigned to α-helical or polyproline-II conformation using DSSP (v4).

To compare α-helical or polyproline-II content in IDPs belonging to different organisms or with different Gene Ontology (GO) terms, we calculated an average percentage of residues assigned to α-helical or polyproline-II conformation by DSSP in the generated conformational ensemble of each IDP. Subsequently, we calculated and compared the average of these average percentages across all IDPs belonging or not belonging to a particular group (e.g. viruses vs all organisms except viruses). To estimate if the difference was significant between two groups (e.g. viruses vs not viruses or archaea+bacteria vs eukaryota), we performed a one-sided Brunner-Munzel test, with the null hypothesis that it was as probable to pick up an IDP with more α-helical or polyproline-II average content from one of these groups as from another group. p-values were calculated using t-distribution as implemented in SciPy v1.16 (scipy.stats.brunnermunzel). To estimate the effect sizes we calculated Cohen’s *d* value ^34^ with the standard error estimated as the standard deviation using the bootstrap procedure with 100000 re-samplings. For each comparison performed using the Brunner-Munzel test, we reported population sizes, mean and standard error of the mean values, p-values, Cohen’s *d* values and degrees of freedom in Extended Data Table 4. We also performed 100000 bootstraps to verify the significance of Brunner-Munzel’s p-values and obtained values similar to the original ones. For the bootstrap procedure, two samples of the size of the original populations were drawn with replacement from a pooled population, as proposed previously ^35^, and Brunner-Munzel’s W test statistics values were used.

Taxon IDs for each IDP/disordered region were obtained directly from UniProt. To group NCBI taxa for different IDPs (e.g. all human IDPs, all eukaryotic IDPs), we used the Environment for Tree Exploration (ETE) framework (version 3) for Python ^36^. Specific GO terms for each IDP/R were obtained directly from UniProt. To access shared (parent, or ascendant) GO terms from specific terms associated with IDPs (i.e. lipid binding, nucleic acid binding, protein binding descend from parent ‘binding’ term) and group together IDPs according to the shared terms (e.g. all ‘binding’-associated IDPs), we downloaded the basic GO version from the Open Biological and Biomedical Ontology (OBO) Foundry (http://purl.obolibrary.org/obo/go/go-basic.obo), containing GO terms and their connections in a form of a graph, which we analyzed using NetworkX ^37^. We combined all GO terms associated with all analyzed DisProt IDPs and all their parent terms, and retained only terms which were connected to at least 50 different IDPs. Obtained GO term IDP groups were analyzed for their α-helical or polyproline-II conformational content as described above.

