## Extended Data for "Decoding conformational heterogeneity across disordered proteomes"

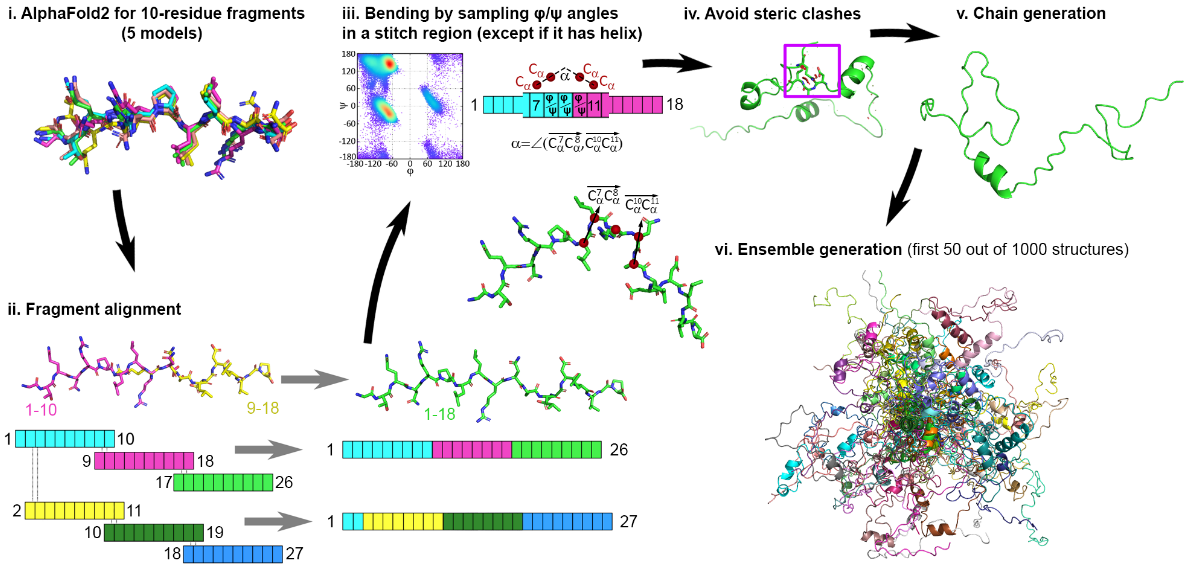


**Extended Data Figure 1.** Schematic representation of AI-IDP. **i.** Fragment generation (5 structures per fragment). **ii.** Fragment alignment with different starting points. **iii.** Flexible physical assembly that preserves the context of local dihedral distributions including an upper limit of the angle formed by pairs of $C_{\alpha}$ atoms (see also Extended Data Table 1). **iv.** Discarding structures with steric clashes (heavy atoms at or closer than 1.18 Å). **v.** One generated chain. **vi.** Ensemble of generated chains.

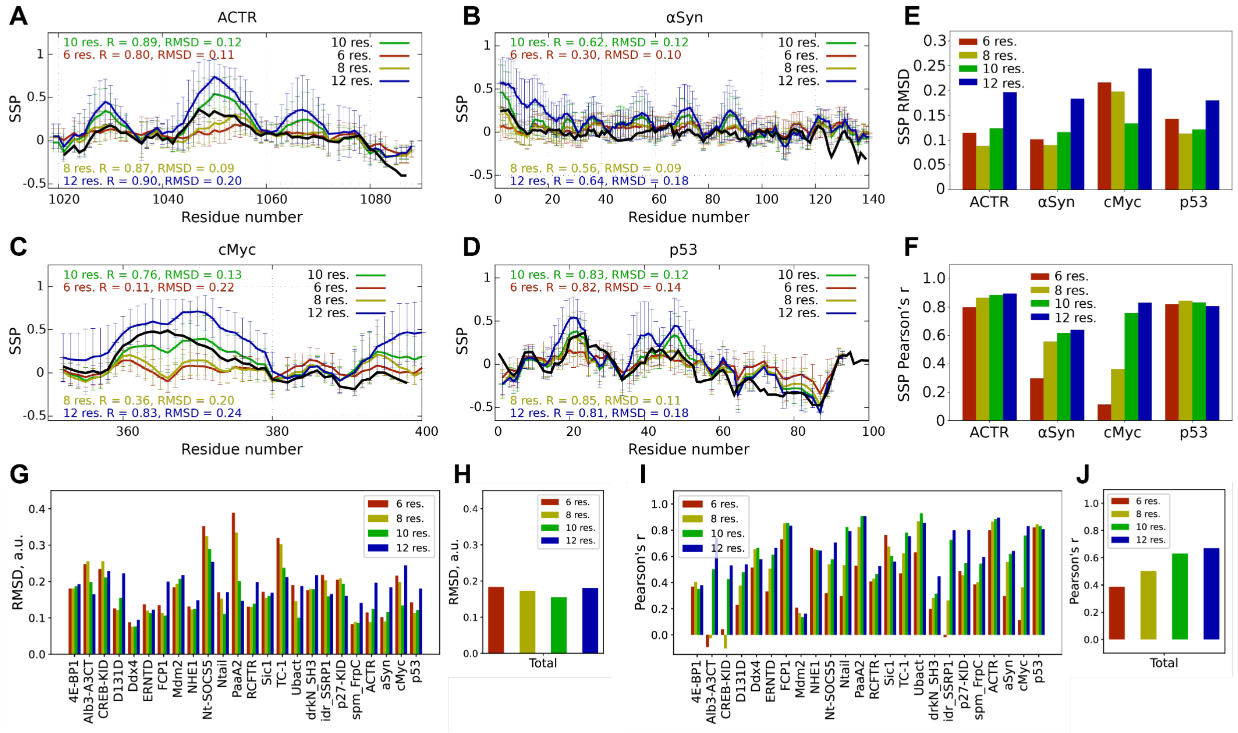

**Extended Data Figure 2.** Influence of fragment size on local structural properties of AI-IDP ensembles. **A-D.** Residue-specific SSP scores derived from experimental chemical shifts (black) and chemical shifts predicted from AI-IDP ensembles generated with 6, 8, 10 and 12-residue fragments for the disordered regions of ACTR (A), α-synuclein (B), cMyc (C) and p53 (D). **E-F.** RMSDs (E) and Pearson correlation coefficients (F) between AI-IDP derived and experimental SSP scores for ensembles generated with 6, 8, 10 and 12-residue fragments. **G-J.** RMSDs (G) and Pearson correlation coefficients (I) between AI-IDP derived and experimental SSP scores for ensembles generated with 6, 8, 10 and 12-residue fragments separately for 24 IDPs, and total RMSDs (H) and correlation coefficients (J) for aggregated data from 24 IDPs.

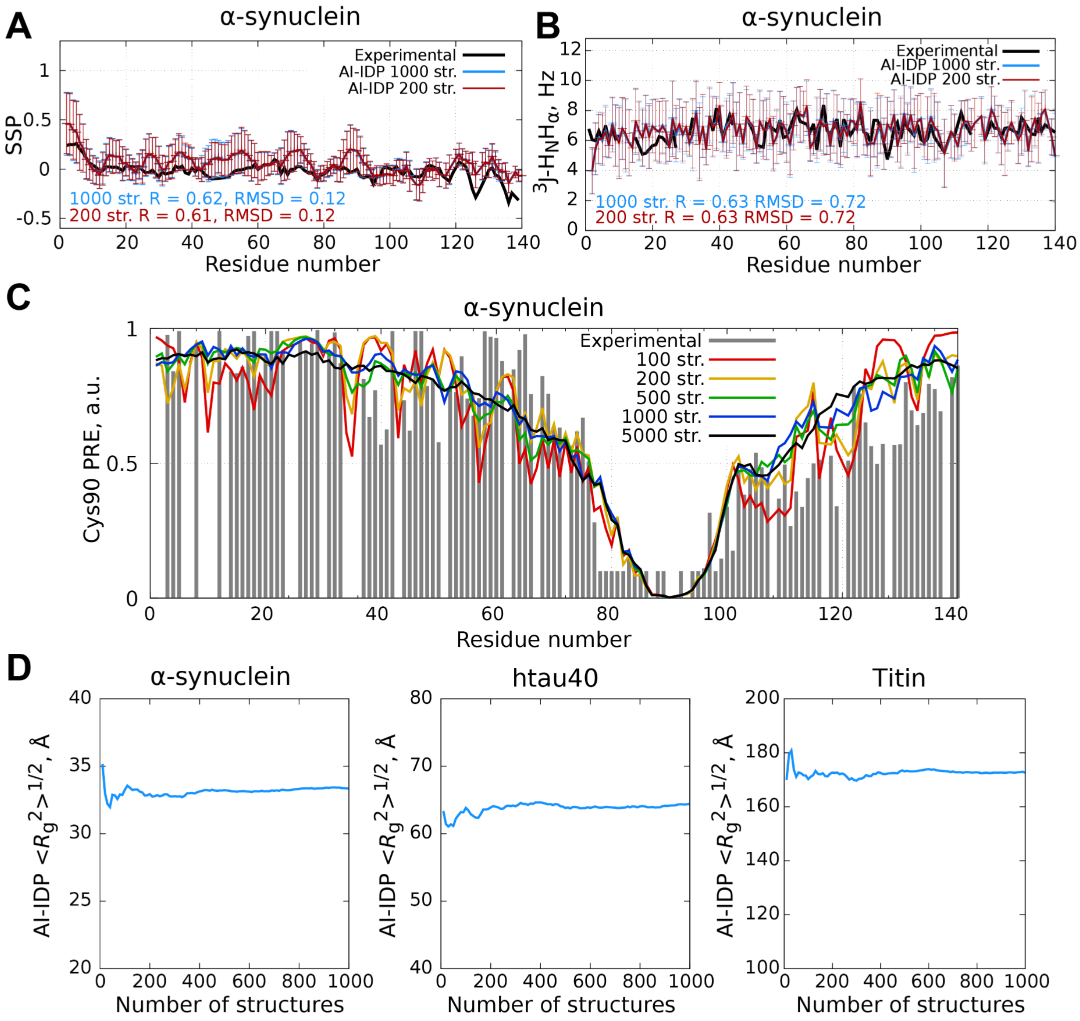


**Extended Data Figure 3.** Influence of AI-IDP ensemble size on back-calculated structural properties. **A-B.** Residue-specific SSP scores derived from experimental chemical shifts (black) and chemical shifts predicted from AI-IDP ensembles of α-synuclein with 1000 (blue line) and 200 (red line) structures. **B.** Experimental $^3J(H_NH_\alpha)$ couplings (black) and couplings calculated from AI-IDP ensembles of α-synuclein with 1000 (blue line) and 200 (red line) structures. **C.** Experimental (black) and AI-IDP-predicted PRE profiles for α-synuclein with a spin label at Cys90 for ensembles of 100 (red), 200 (yellow), 500 (green), 1000 (blue) and 5000 structures (black line). **D.** Root mean square radius of gyration ($R_g$) from AI-IDP ensembles for 140-residue α-synuclein, 441-residue htau40 and 2152-residue Titin for increasing number of ensemble structures.

Together, the analyses demonstrate that approximately 100-200 AI-IDP structures (for SSP scores and $^3J(H_NH_\alpha)$ couplings) and around 500-1000 structures (for PREs and $R_g$) are required for the convergence of ensemble averages of back-calculated structural parameters.

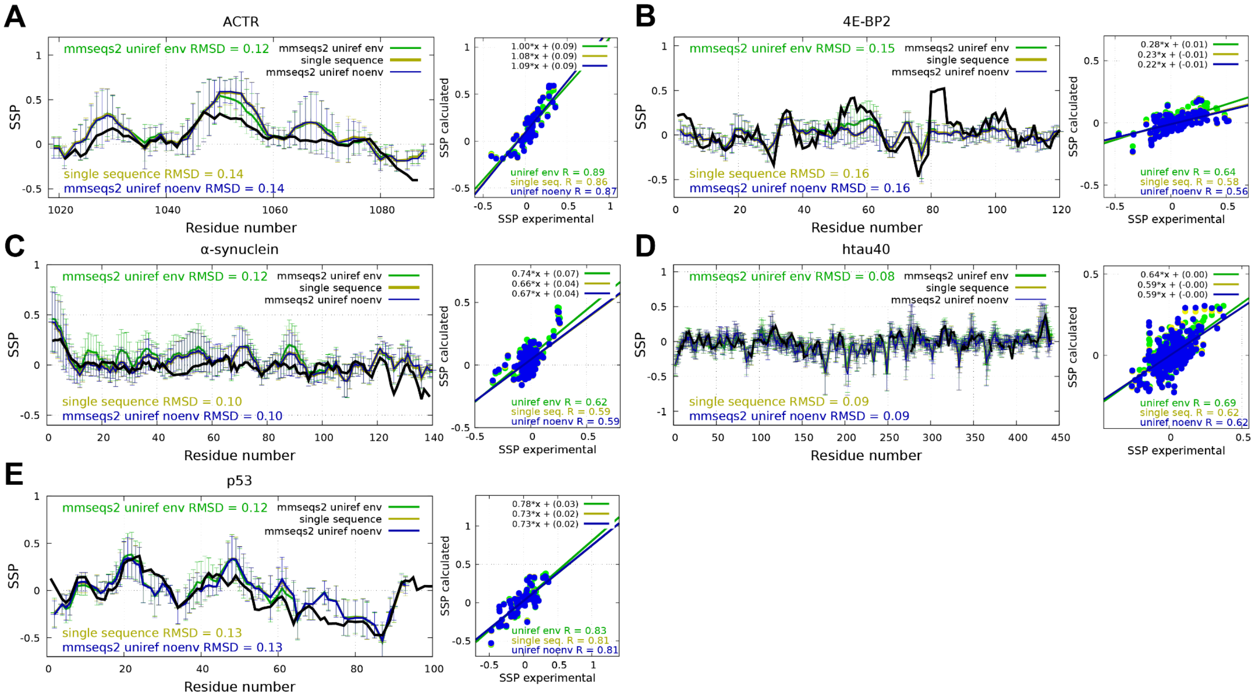


**Extended Data Figure 4.** Influence of sequence alignment mode (MSA with UniRef and environmental databases, UniRef only or single sequence) on AI-IDP ensembles assessed by SSP. **A-E.** Sequence and correlation plots of SSP scores derived from experimental NMR chemical shifts (black) and chemical shifts predicted from AI-IDP ensembles of the disordered regions of ACTR (A), 4E-BP2 (B), α-synuclein (C), htau40 (D) and p53 (E) using fragments generated by AlphaFold2 with different sequence alignment modes. RMSDs are shown on residue-specific SSP score plots, Pearson correlation coefficients between AI-IDP and experimental SSPs on correlation plots.

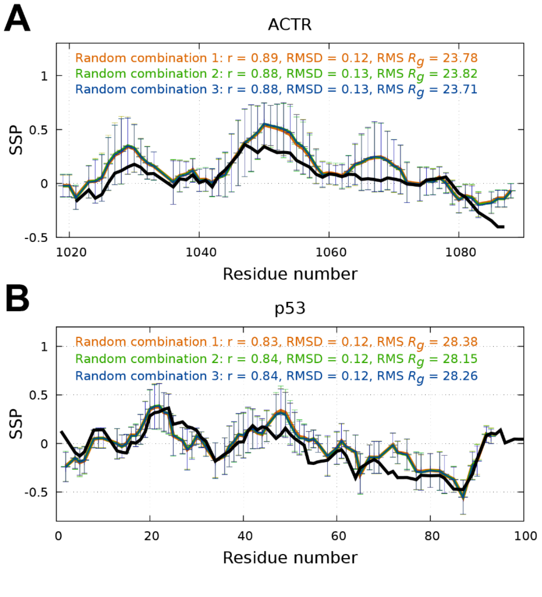


**Extended Data Figure 5.** Influence of different random combinations of fragment structures and residue-specific ϕ/ψ angle pairs on SSP scores derived from AI-IDP ensembles. **A,B.** Residue-specific SSP scores derived from experimental chemical shifts (black) and shifts calculated from AI-IDP ensembles of the disordered regions of ACTR (A) and p53 (B). RMSDs, Pearson correlation coefficients between AI-IDP and experimental SSP scores as well as root mean squared radii of gyration are indicated.

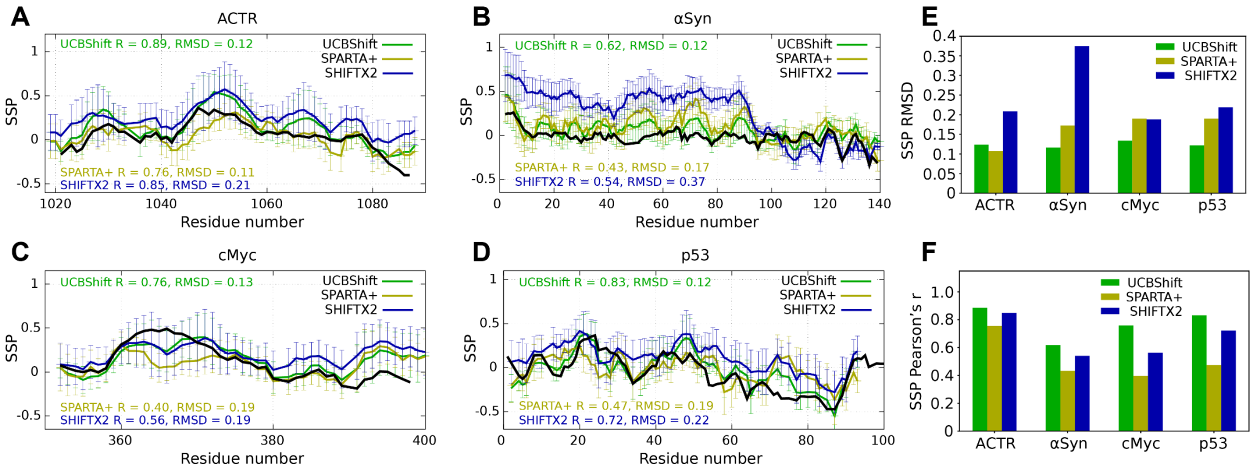


**Extended Data Figure 6.** Influence of chemical shift predictors on SSP scores derived from AI-IDP ensembles. **A-D.** Residue-specific SSP scores derived from experimental chemical shifts (black) and shifts calculated with UCBShift, SPARTA+ and SHIFTX2 from the AI-IDP ensembles of the disordered regions of ACTR (A), α-synuclein (B), cMyc (C) and p53 (D). **E,F.** RMSDs (E) and Pearson correlation coefficients (F) between AI-IDP-derived and experimental SSP scores for different chemical shifts predictors.

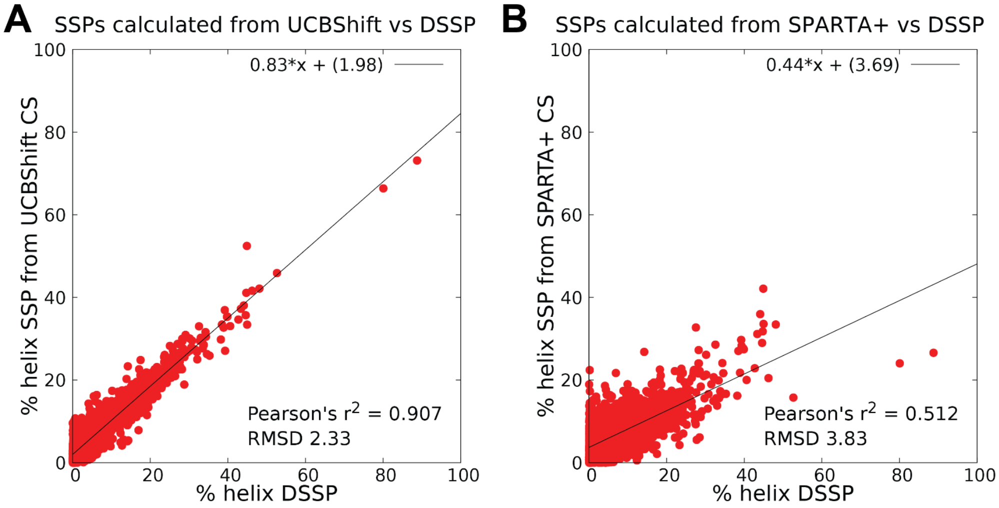


**Extended Data Figure 7.** Accuracy and precision of the transient α-helical content determination from SSP scores using UCBShift and SPARTA+ as chemical shift predictors. **A,B.** Transient α-helical content was determined using DSSP algorithm in IDP structures generated by AI-IDP and compared to the sequence average of positive-only SSP scores calculated from chemical shifts predicted using UCBShift (A) and SPARTA+ (B). Pearson correlation values, RMSDs, regression line and linear fit equation are shown. Comparison was performed on all 3389 IDPs in DisProt for which AI-IDP ensembles were generated.

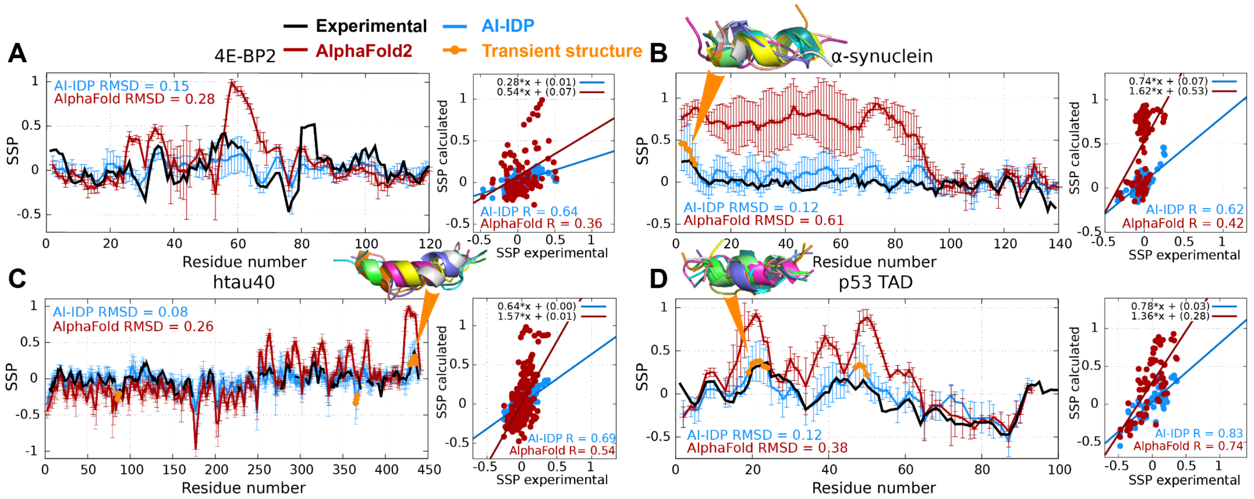


**Extended Data Figure 8.** Local structural properties derived from AI-IDP ensembles, AlphaFold2 structures and from experimental NMR chemical shifts. **A-D.** Sequence and correlation plots of SSP scores derived from the AI-IDP ensembles, AlphaFold2 structures and experimental NMR chemical shifts of the proteins 4E-BP2 (A), α-synuclein (B), htau40 (C) and p53 transactivation domain (D). Residues with transient α-helical structure are marked in orange; corresponding AI-IDP ensemble structures are shown above. RMSDs are shown on residue-specific SSP score plots, Pearson correlation coefficients between AI-IDP and experimental SSPs on correlation plots.

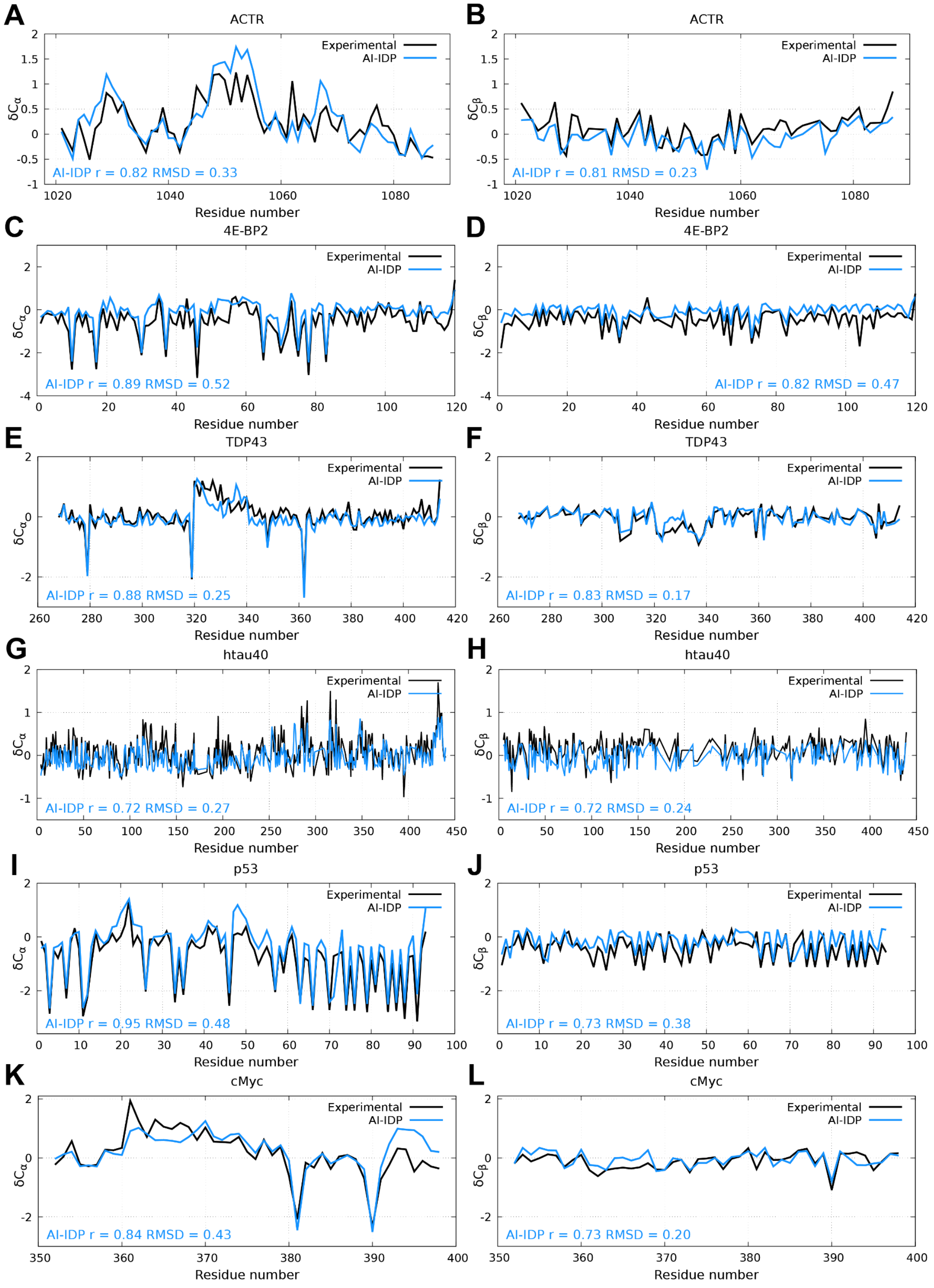


**Extended Data Figure 9.** Chemical shifts derived from AI-IDP ensembles closely match experimental values. **A-L.** Residue-specific secondary $C_\alpha$, $C_\beta$ chemical shifts derived from experimental NMR chemical shifts (black) and shifts back-calculated from AI-IDP ensembles (blue lines) for ACTR (A,B), 4E-BP2(C,D), TDP43 (E,F), htau40 (G,H), p53 (I,J) and cMyc (K,L). Secondary chemicals shifts are calculated as the difference between $C_\alpha$, $C_\beta$ chemical shifts and RefDB residue-specific random coil values (Zhang et al. 2003, *J. Biomol. NMR* 25:173-195).

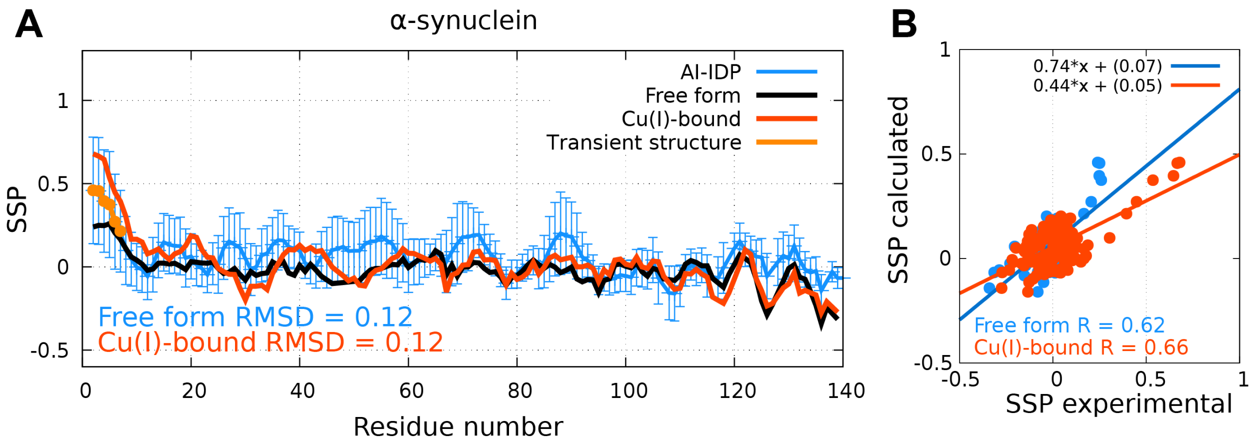


**Extended Data Figure 10.** Sensitivity of the local structure in α-synuclein to Cu(I)-binding. **A.** Residue-specific SSP scores derived from the experimental NMR chemical shifts observed in α-synuclein in the absence (black) and presence of Cu(I) (red), as well as SSP scores derived from α-synuclein's AI-IDP ensemble (blue). In both cases, synuclein was in MES 20 mM, NaCl 100 mM, pH 6.5, 288 K. Residues with transient α-helical structure in the AlphaFold-IDP ensemble are marked in light orange. Error bars represent the standard deviation of the SSP values from 1000 structures. RMSD values between AI-IDP-derived SSP scores and SSP scores derived from experimental chemical shifts are shown. **B.** Correlation plot between SSP scores derived from the experimental NMR chemical shifts of α-synuclein in the absence and presence of Cu(I) and those derived from α-synuclein's AI-IDP ensemble. The respective Pearson correlation coefficients are displayed.

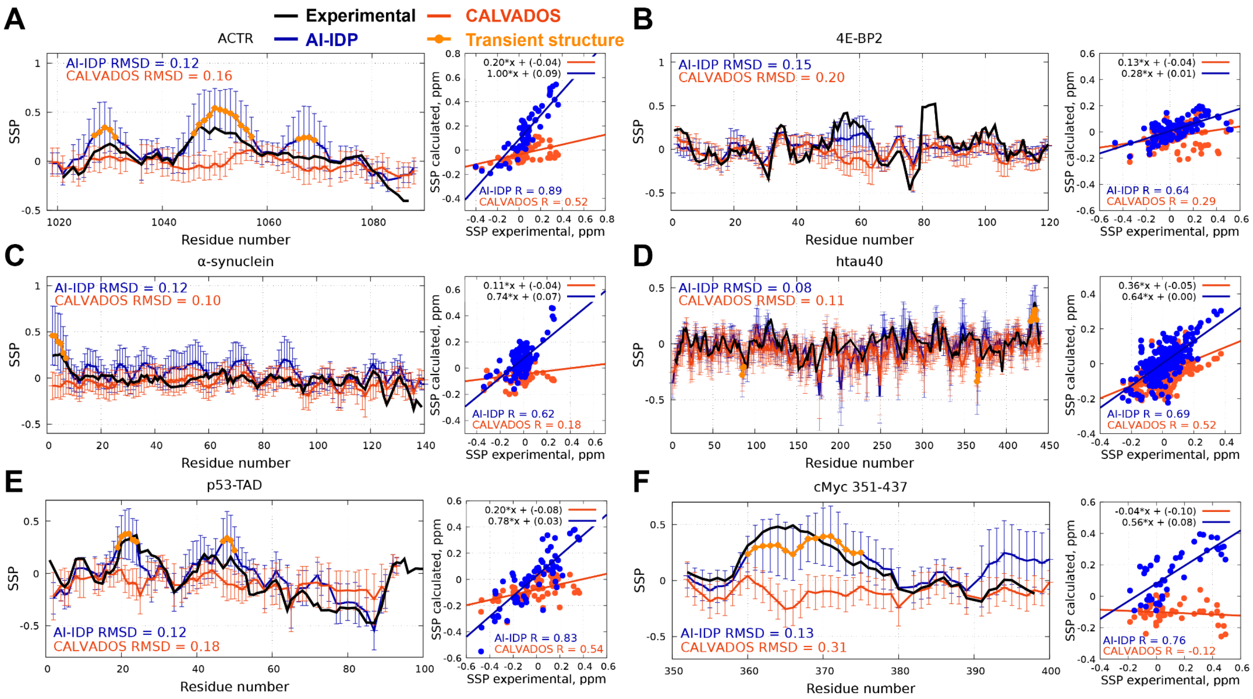


**Extended Data Figure 11.** Transient secondary structures in disordered proteins are captured by AI-IDP but not by CALVADOS. **A-F.** Residue-specific SSP scores and SSP correlation plots derived from AI-IDP and CALVADOS ensembles, as well as experimental data, for ACTR (A), 4E-BP2(B), α-synuclein (C), htau40 (D), p53 (E) and cMyc (F). Residues with transient α-helical or extended (in the case of htau40) structure are marked in orange. Correlation plots display regression line, equation and Pearson correlation coefficient between ensemble-derived and experimental SSPs for both methods.

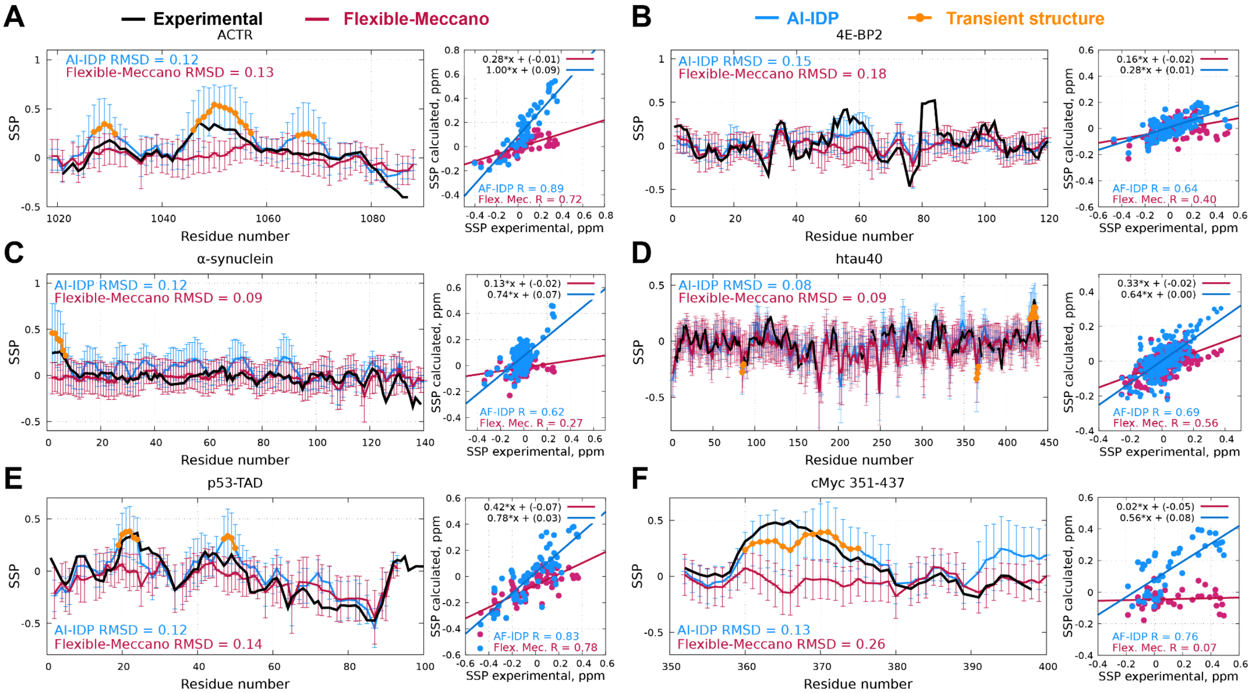

**Extended Data Figure 12.** Transient secondary structures in disordered proteins are captured by AI-IDP but not by Flexible-Meccano. **A-F.** Residue-specific SSP scores and SSP correlation plots derived from AI-IDP and Flexible-Meccano ensembles, as well as experimental data, for ACTR (A), 4E-BP2(B), α-synuclein (C), htau40 (D), p53 (E) and cMyc (F). Residues with transient α-helical or extended (in the case of htau40) structure are marked in orange. Correlation plots display regression line, equation and Pearson correlation coefficient between ensemble-derived and experimental SSPs for both methods.

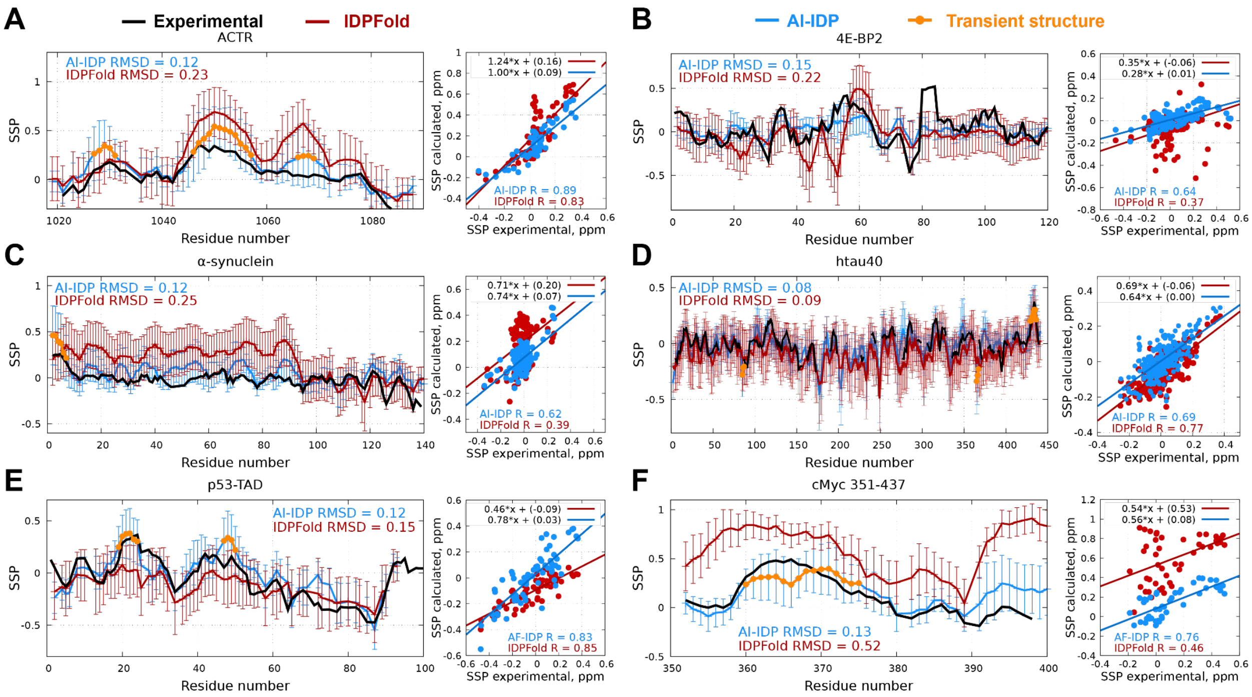


**Extended Data Figure 13.** Transient secondary structures in disordered proteins are captured better by AI-IDP. **A-F.** Residue-specific SSP scores and SSP correlation plots derived from AI-IDP and IDPFold ensembles, as well as experimental data, for ACTR (A), 4E-BP2(B), α-synuclein (C), htau40 (D), p53 (E) and cMyc (F). Residues with transient α-helical or extended (in the case of htau40) structure are marked in orange. Correlation plots display regression line, equation and Pearson correlation coefficient between ensemble-derived and experimental SSPs for both methods.

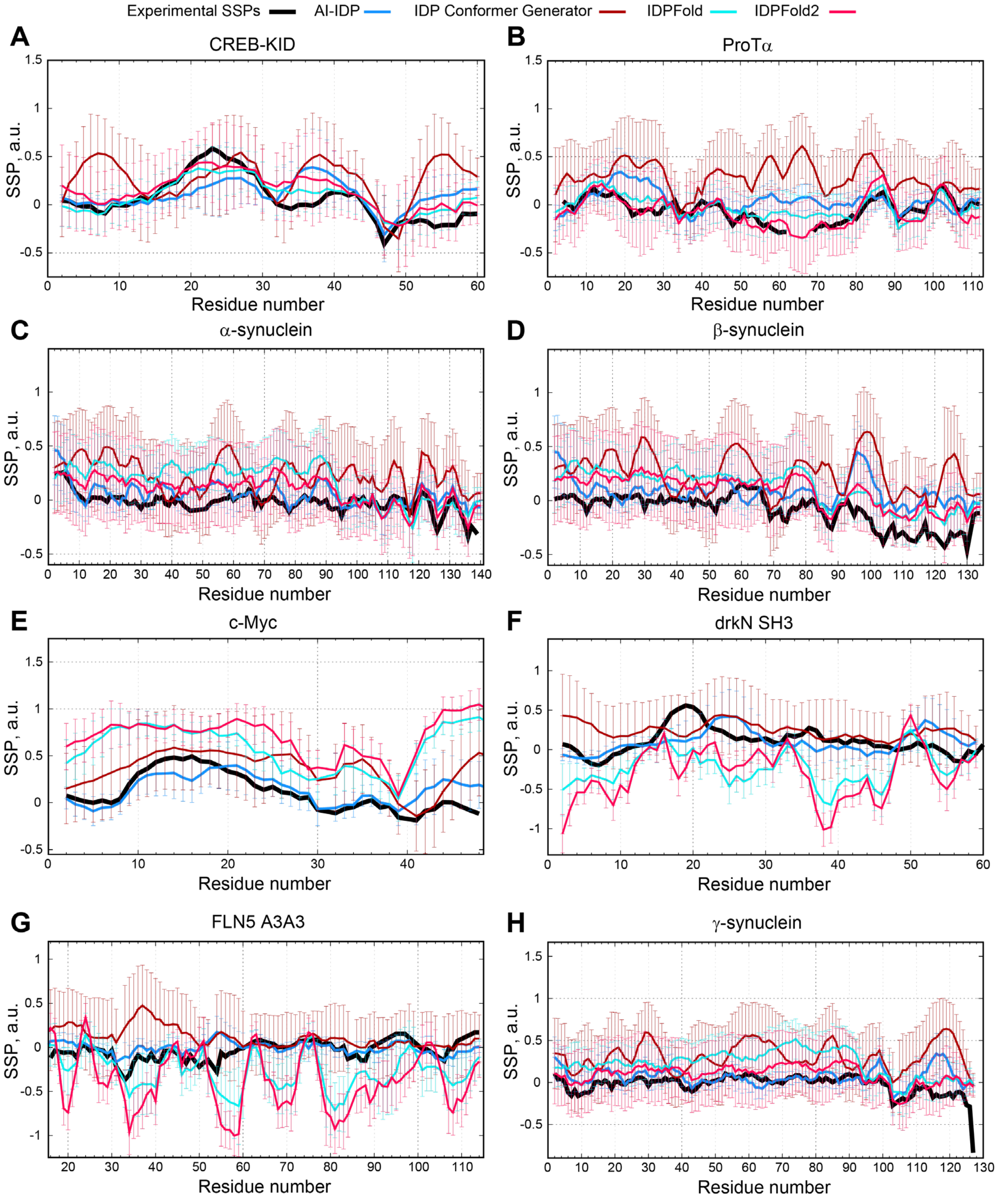


**Extended Data Figure 14.** IDPFold and IDPFold2 capture transient secondary structure well in some IDPs (A,B) but spuriously detect it in others (C-F). **A-H.** Residue-specific SSP scores derived from AI-IDP, IDPFold, IDPFold2 and IDP Conformer Generator ensembles, as well as experimental data, for CREB-KID (A), ProTα (B), α-synuclein (C), β-synuclein (D), c-Myc (E), drkN SH3 domain (F), FLN5 A3A3 variant (G) and γ-synuclein (H).

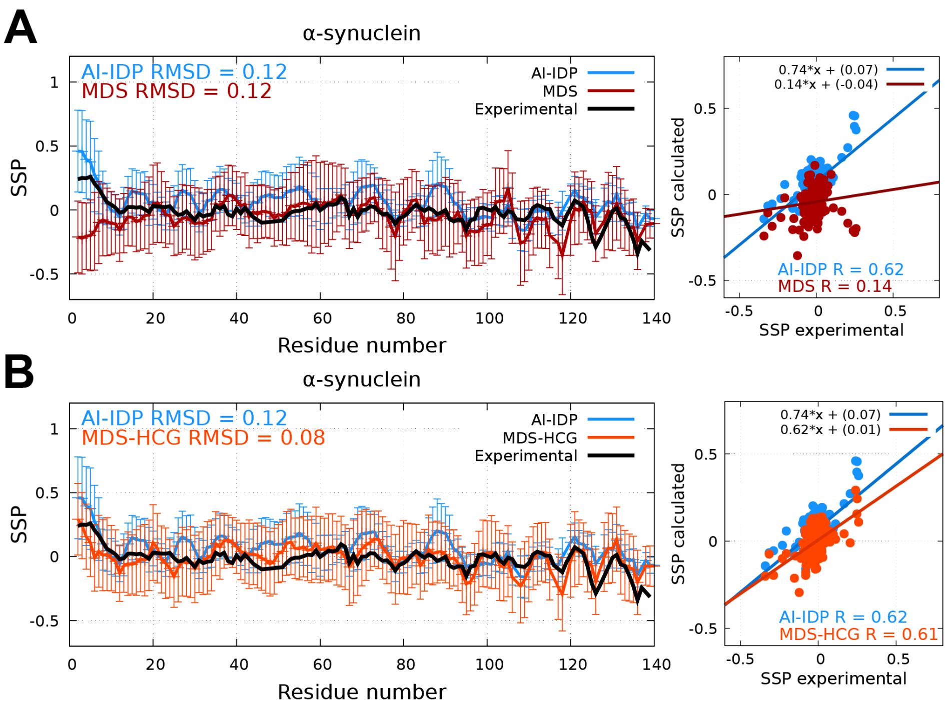


**Extended Data Figure 15.** Accuracy of local structural properties in conformational ensembles generated by AI-IDP and molecular dynamics simulations. **A, B.** Residue-specific SSP scores and correlation plots derived from NMR chemical shifts back-calculated from AI-IDP (blue) and MDS (magenta; A) or MDS-HCG (orange; B) ensembles versus SSP scores derived from experimental NMR chemical shifts (black). Correlation plots display regression line, equation and Pearson correlation coefficient between ensemble-derived and experimental SSPs.

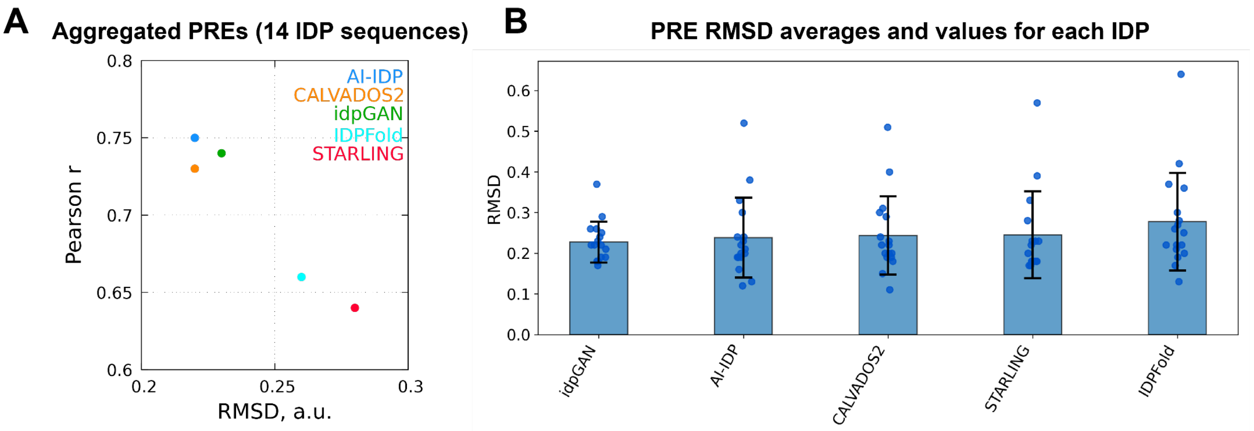


**Extended Data Figure 16.** Medium-range order observed in generated ensembles. **A.** Accuracy of PRE predictions in AI-IDP (light blue), CALVADOS2 (orange), idpGan (green), IDPFold (cyan) and STARLING (magenta) ensembles. Each dot represents a comparison between ensemble-derived and experimental PREs for aggregated PREs for 14 IDPs (Extended Data Table 4). **B.** Accuracy of PRE predictions as measured by IDP-specific and average RMSDs between ensemble-derived and experimental PREs. AI-IDP achieves second-lowest average PRE RMSDs, yielding only to idpGAN.

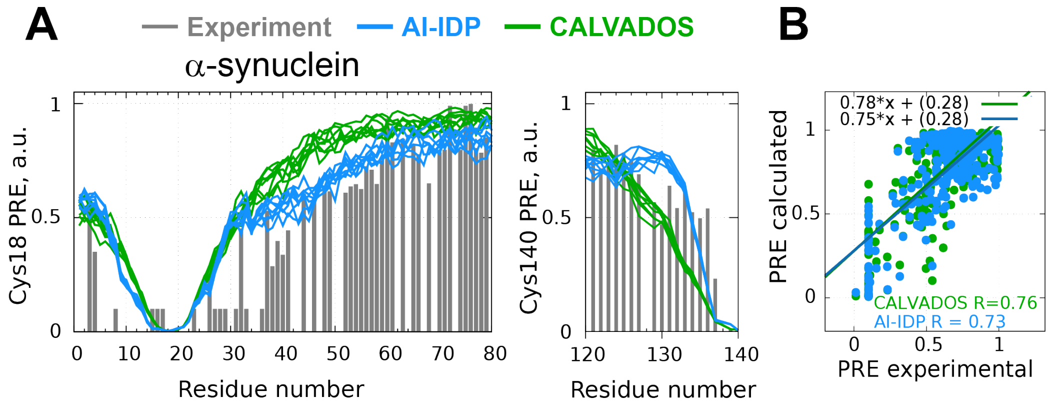


**Extended Data Figure 17.** Medium-range order observed in AI-IDP and CALVADOS ensembles. **A.** Experimental and AI-IDP or CALVADOS-derived PRE profiles of α-synuclein (spin label at Cys18 or Cys140). CALVADOS PRE profiles lack the "shoulder" between residues 30-45 that is present in AI-IDP-derived and experimentally-determined PRE profiles. In addition, the PRE-broadening downstream of the Cys140 spin label position is broader in case of CALVADOS when compared to either the experimental or the AI-IDP-derived PRE profile. Five different 1000-residue AI-IDP or CALVADOS ensembles were used for PRE predictions, shown as five distinct blue lines, to demonstrate reproducibility. **B.** Correlation between experimental and predicted PRE intensity ratios for all α-synuclein spin label positions (Cys18, Cys90, Cys140) together.

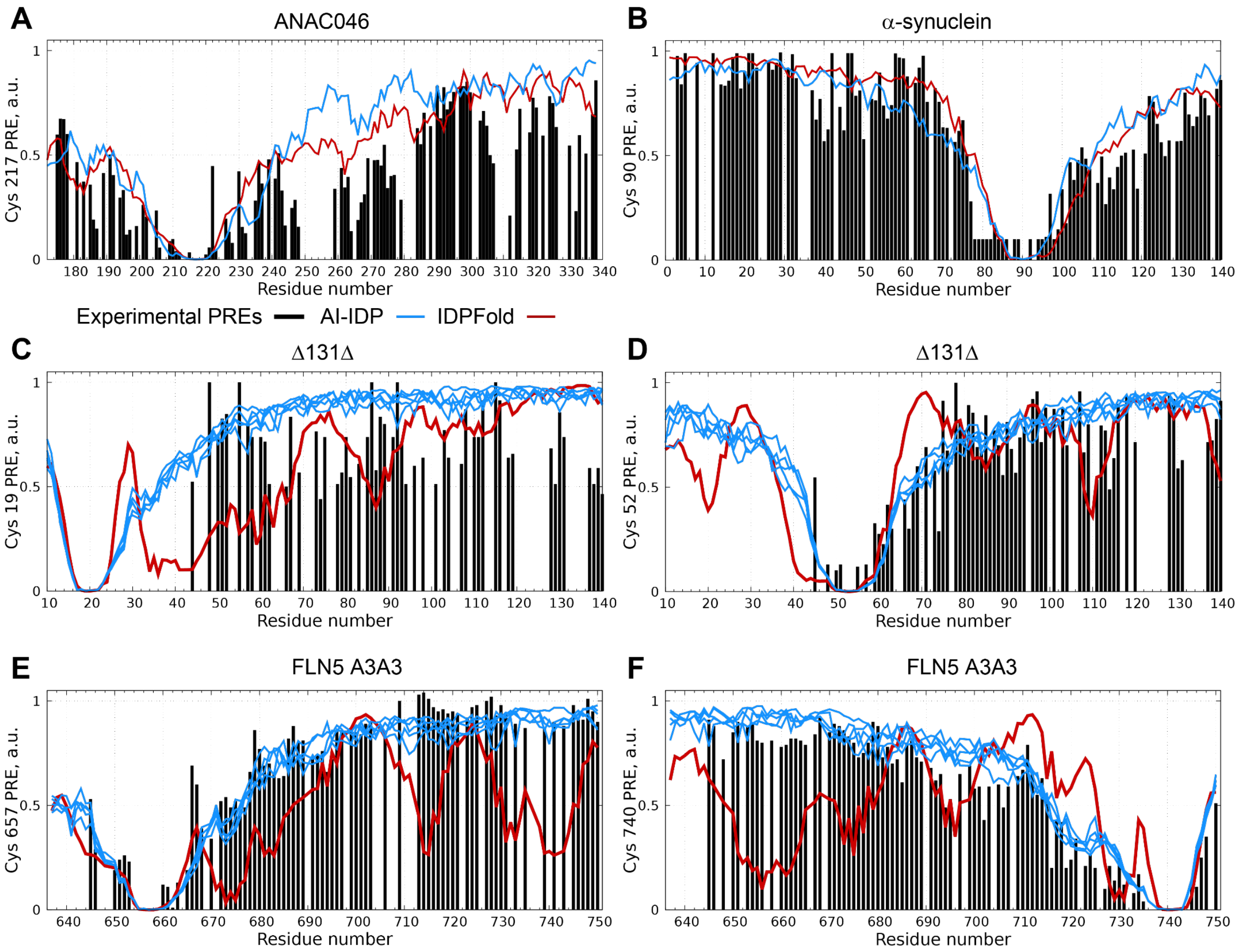


**Extended Data Figure 18.** Medium-range order observed in AI-IDP and IDPFold2 ensembles. **A, B.** Experimental and AI-IDP or IDPFold-derived PRE profiles of ANAC046 (spin label at Cys217) and α-synuclein (spin label at Cys90). IDPFold PRE profiles match experimental profiles well. **C-F.** Experimental and AI-IDP or IDPFold-derived PRE profiles of Δ131Δ fragment of staphylococcal nuclease (spin label at Cys19 and Cys52) and FLN5 A3A3 variant (spin label at Cys657 and Cys740). IDPFold PRE profiles feature multiple spurious contacts across IDP sequence.

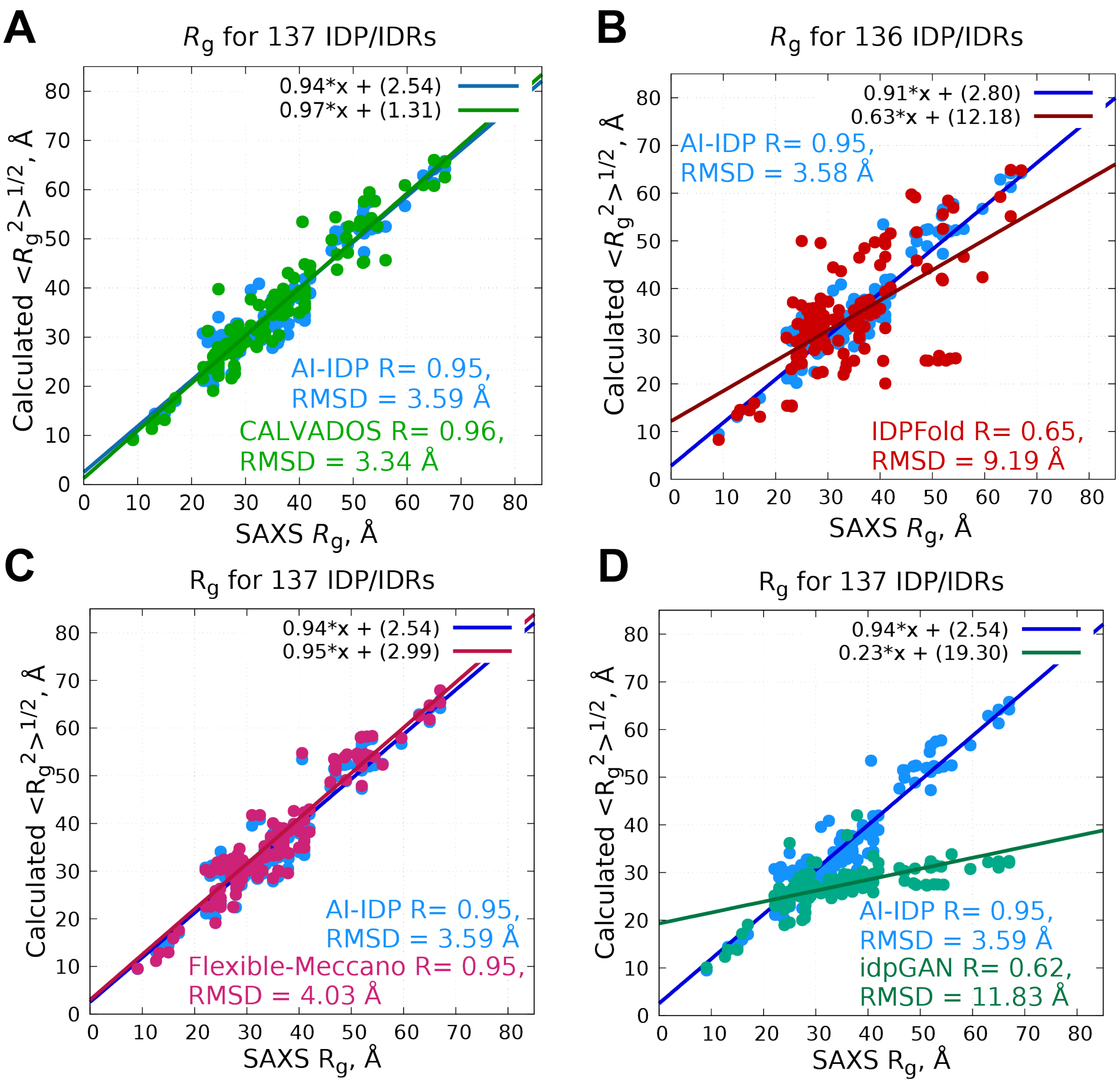


**Extended Data Figure 19.** Global dimensions observed in AI-IDP and CALVADOS ensembles. **A.** Correlation plot of AI-IDP-derived and CALVADOS-derived $R_g$ values versus $R_g$ values obtained from SAXS measurements for 137 IDPs. **B.** Correlation plot of AI-IDP-derived and IDPFold-derived $R_g$ values versus $R_g$ values obtained from SAXS measurements for 136 IDPs (excluding 467-residue MAP2c protein). **C.** Correlation plot of AI-IDP-derived and idpGAN-derived $R_g$ values versus $R_g$ values obtained from SAXS measurements for 137 IDPs. AI-IDP and CALVADOS2 correctly capture global dimensions of IDPs, whereas IDPFold and idpGAN are not.

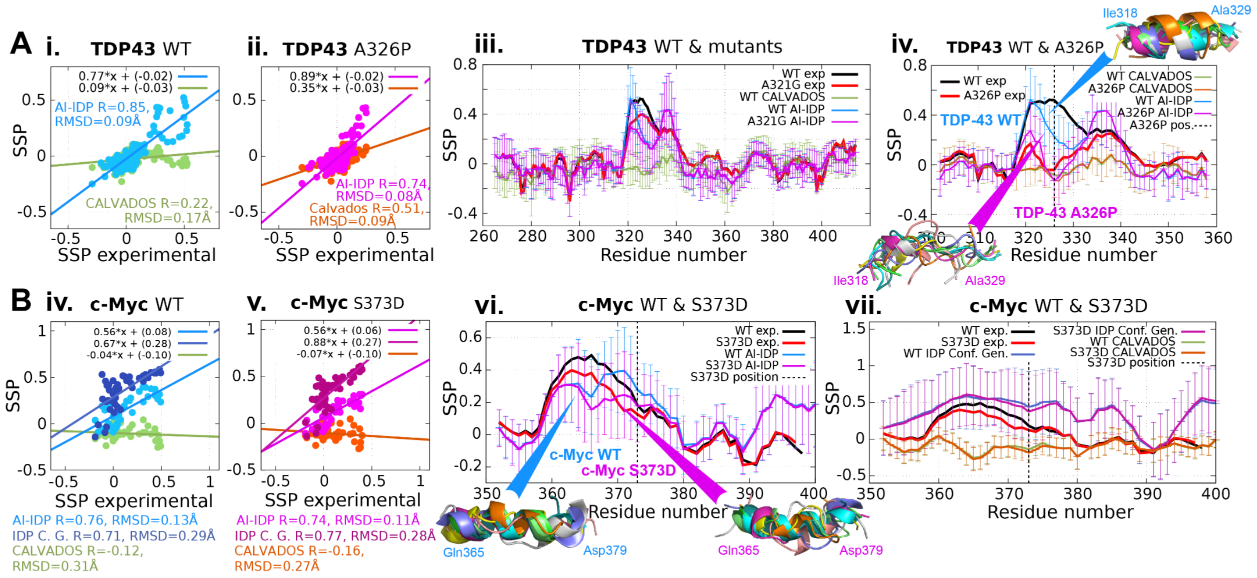


**Extended Data Figure 20.** AI-IDP captures the impact of point mutations on transient secondary structure in IDPs. **A. TDP-43.** (i, ii) Correlation and RMSD between residue-specific SSP scores derived from AI-IDP and CALVADOS ensembles and SSP scores derived from experimental NMR chemical shifts for the C-terminal domain (residues 267–414) of wild-type TDP-43 and its A326P mutant. Pearson correlation coefficients (R), RMSD values, regression lines and linear fit equations are shown. (iii, iv) Residue-specific SSP scores derived from AI-IDP and CALVADOS ensembles, and those expermentially determined for the C-terminal domain of wild-type, A321G, and A326P TDP-43. The mutant proteins show less transient helicity for residues 310-350 both experimentally and in the AI-IDP, but not the CALVADOS ensemble. **B, c-Myc.** (iv, v) Correlation and RMSD between SSP scores derived from AI-IDP, CALVADOS and IDPConformerGenerator ensembles for wild-type and phosphomimetic S373D-mutant c-Myc. (vi, vii) Comparison of AI-IDP-, CALVADOS-, IDPConformerGenerator-derived and experimental SSP profiles for wild-type and S373D-mutant c-Myc, showing loss of transient helicity for residues 350-380. Representative AI-IDP conformers are shown.

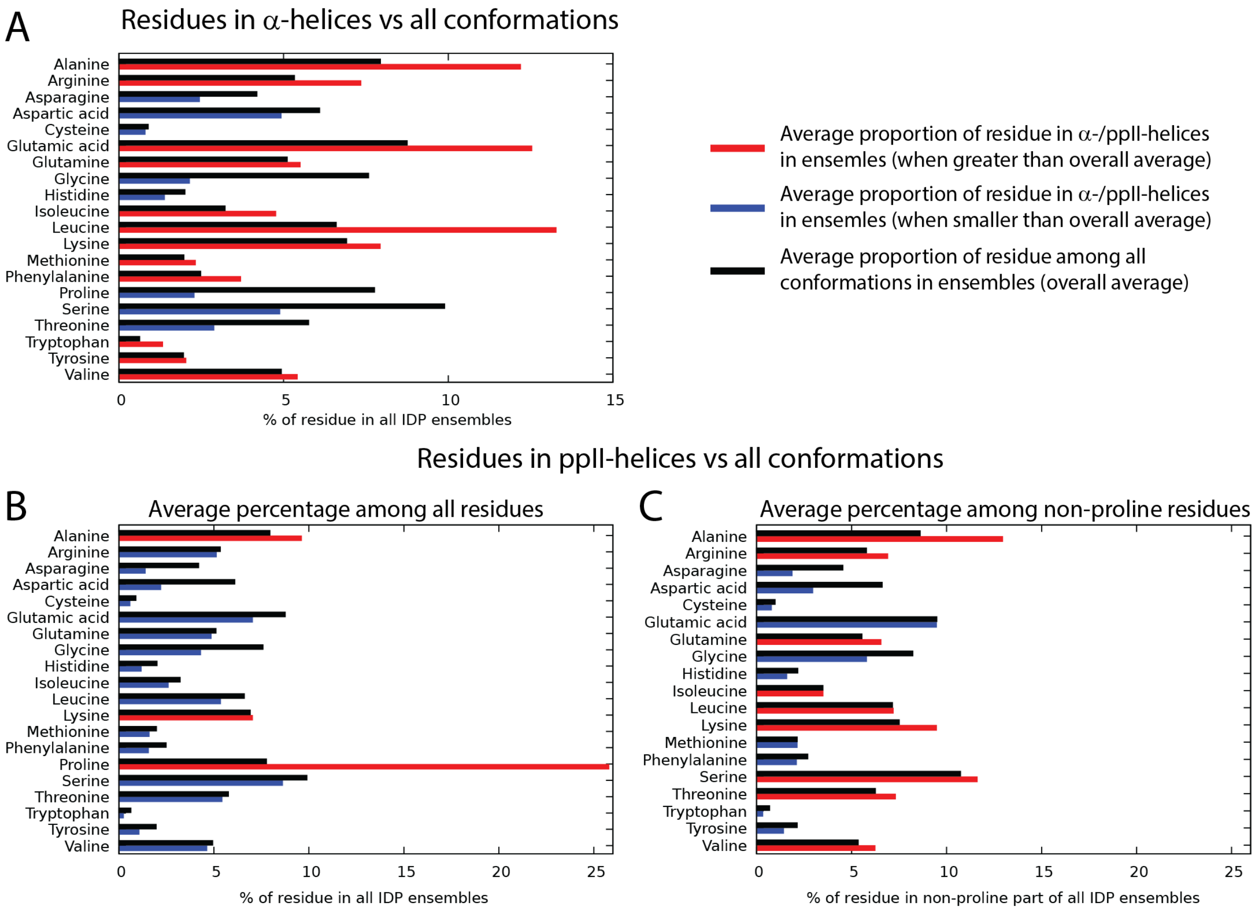


**Extended Data Figure 21.** Prevalence of amino acid types in secondary structure elements in AI-IDP ensembles. Alanine, arginine, leucine, lysine, glutamic acid, isoleucine residues are more prevalent in α-helices, whereas glycine, proline, aspartic acid and threonine residues are less prevalent. In polyproline-II helices, proline, alanine and serine residues are more prevalent. **A.** Average percentage of amino acid types across all DSSP-identified α-helices in all 1000 AI-IDP structures of all analyzed DisProt IDPs (red, blue bars) versus average percentage of residue types across all full sequences of all DisProt IDPs (black bars). **B.** Average percentage of residue types across all DSSP-identified polyproline-II helices in all 1000 AI-IDP structures of all analyzed DisProt IDPs (red, blue bars) versus average percentage of residue types across all full sequences of all DisProt IDPs (black bars). **C.** Same as (B) but proline residues removed from polyproline-II helix sequences and from full sequences for averaging purposes.

**Extended Data Table 1.** Optimization of the AI-IDP curvature angle for the flexible assembly of fragments. Different limiting angles between $C_\alpha^{i}$, $C_\alpha^{i+1}$…$C_\alpha^{i+3}$, $C_\alpha^{i+4}$, in the juncture points of fragments were tested on a randomly chosen subset of ~1/3 (43) out of 137 IDPs/IDRs with known experimentally determined $R_g$ to find a minimal RMSD between calculated $<R_g^2>^{1/2}$ from AI-IDP ensembles and experimentally determined $R_g$. Proteins in the Extended Data Table 2 were excluded from this subset. Pearson correlation coefficients and RMSD values are reported for a chosen subset, remaining IDPs/IDRs and all 137 IDPs/IDRs. The optimized angle of 40° is highlighted in bold. Ensembles of 500 conformers were generated for the purpose of $C_\alpha^{i}$, $C_\alpha^{i+1}$…$C_\alpha^{i+3}$, $C_\alpha^{i+4}$ angle optimization, while in Fig. 1 Pearson correlation coefficients and RMSD values obtained with 1000 conformers are reported.

| $C_\alpha^{i}$,$C_\alpha^{i+1}$…$C_\alpha^{i+3}$,$C_\alpha^{i+4}$ value | Subset of 43/137 IDP/Rs | | Remaining 94 IDP/Rs | | All 137 IDP/Rs | |
|---|---|---|---|---|---|---|
| | RMSD, Å | Pearson r | RMSD, Å | Pearson r | RMSD, Å | Pearson r |
| 25° | 3.11 | 0.968 | 3.80 | 0.947 | 3.59 | 0.953 |
| 30° | 3.11 | 0.967 | 3.80 | 0.944 | 3.60 | 0.951 |
| 35° | 3.20 | 0.964 | 3.78 | 0.945 | 3.60 | 0.951 |
| **40°** | **2.98** | **0.968** | **3.84** | **0.944** | **3.60** | **0.951** |
| 45° | 3.23 | 0.963 | 3.84 | 0.945 | 3.61 | 0.951 |
| 50° | 3.25 | 0.963 | 4.00 | 0.943 | 3.64 | 0.950 |
| 55° | 3.39 | 0.960 | 4.08 | 0.943 | 3.67 | 0.949 |
| 60° | 3.53 | 0.962 | 4.22 | 0.943 | 3.72 | 0.949 |
| 70° | 4.14 | 0.954 | 4.63 | 0.941 | 3.81 | 0.947 |
| 80° | 4.66 | 0.954 | 5.22 | 0.939 | 3.95 | 0.945 |
| Comparison: CALVADOS2 (1000 structures) | | | 3.58 | 0.951 | 3.34 | 0.958 |

**Extended Data Table 2.** Experimental conditions of the NMR chemical shifts.

| **Protein** | **Buffer** | **pH** | **Temperature** |
|---|---|---|---|
| 4E-BP2 | NaP 30 mM, NaCl 100 mM | 6.0 | 283 K |
| ACTR | NaP 10 mM, NaCl 50 mM | 6.7 | 304 K |
| α-synuclein | MES 20 mM, NaCl 100 mM | 6.5 | 288 K |
| α-synuclein (Cu(I)) | MES 20 mM, NaCl 100 mM | 6.5 | 288 K |
| BRCA1 | 25 mM MES, 25 mM NaCl,<br>100mM Arginine HCl | 5.5 | 298 K |
| cMyc WT | 50 mM KPi, 500mM $NH_4Cl$, | 6.5 | 277 K |
| cMyc S373D | 50 mM KPi, 500mM $NH_4Cl$, | 6.5 | 277 K |
| p53 | 20 mM MES, 20 mM NaCl, 10 mM TCEP | 6.0 | 298 K |
| TDP-43 WT | 20 mM MES | 6.1 | 283 K |
| TDP-43 A321G | 20 mM MES | 6.1 | 283 K |
| TDP-43 A326P | 20 mM MES | 6.1 | 283 K |
| Tau | NaP 50 mM | 6.8 | 278 K |

**Extended Data Table 3.** Sources of NMR chemical shifts for comparing accuracy of SSPs derived from IDP ensembles generated by different methods (40 IDP/IDRs).

| **Protein** | **Chemical shift source** |
|---|---|
| 4E-BP1 | BMRB 27936 |
| 4E-BP2 | BMRB 19114 |
| Ab40 | BMRB 17796 |
| ACTR | BMRB 15397 |
| Alb3-A3CT | BMRB 25838 |
| ANAC046 | BMRB 51033 |
| Ash1 | BMRB 26719 |
| Alpha-synuclein | Miotto et al., *J. Am. Chem. Soc.* 137, 6444–6447 (2015) |
| Beta-synuclein | BMRB 15298 |
| c-Myc | BMRB 27414 |
| CREB-KID | BMRB 6784 |
| D131D | BMRB 52367 |
| Ddx4 | BMRB 51711 |
| drkN_SH3 | BMRB 51327 |
| ERNTD | BMRB 51972 |
| FCP1 | BMRB 16296 |
| FLN5 A3A3 | Streit et al, *Nature*, 633, 232–239 (2024) |
| Gamma-synuclein | BMRB 7244 |
| hnRNPA2-LC_266-341 | BMRB 27649 |
| hnRNPA2-LC_WT | BMRB 27123 |
| htau40 | Ukmar-Godec et al., *Sci. Adv.* 6, eaba3916 (2020) |
| HTLV-1_HBZ | BMRB 27517 |
| idr_SSRP1 | BMRB 11511 |
| KISS-1 | BMRB 26935 |
| Mdm2 | BMRB 28011 |
| NHE1 | BMRB 26755 |
| Nsp2_CtIDR | BMRB 50687 |
| Ntail | Gely et al., *J. Mol. Recognit.* 23, 435–447 (2010) |
| Nt-SOCS5 | BMRB 19966 |
| OPN | BMRB 15519 |
| p15PAF | BMRB 19332 |
| p27-KID | BMRB 6112 |
| p53-TAD | BMRB 51984 |
| PaaA2 | BMRB 18841 |
| ProTalpha | BMRB 27215 |
| RCFTR | BMRB 15336 |
| Sic1 | BMRB 16657 |
| spm_FrpC | BMRB 26530 |
| TC-1 | BMRB 15141 |
| Ubact | BMRB 51116 |

**Extended Data Table 4.** Sources of NMR chemical shifts for comparing accuracy of SSPs derived from IDP ensembles generated by different methods (40 IDP/IDRs).

| **Protein** | **Chemical shift source** |
|---|---|
| ACTR | Iešmantavičius et al, *J. Am. Chem. Soc.* 135 (27), 10155–10163 (2013) |
| ANAC046 | Due et al., *Prot. Sci.* 34 (6), p.e70142 (2025) |
| D131D | Gillespie et al, *J. Mol. Biol.* 268 (1), 158–169 (1997) |
| ERNTD | Du et al., *Nature* 638, 1130–1138 (2025) |
| FLN5A3A3 | Streit et al., *Nature* 633, 232–239 (2024) |
| FUS LC | Ryan et al., *Mol. Cell* 69, 465–479.e7 (2018), Figure S1 |
| OPN | Konrat et al., *J. Magn. Reson.* 241, 74–85 (2014) |
| Pdx1-C | Cook et al., *J. Phys. Chem. B* 123 (1), 106–116 (2018) |
| Sic1 | Mittag et al., *Structure* 18 (4), 494–506 (2010) |
| Sml1 | Danielsson et al., *Biochemistry* 47 (50), 13428–13437 (2008) |
| Alpha-synuclein | Bertoncini et al., *Proc. Natl. Acad. Sci.* 102, 1430–1435 (2005) |

**Extended Data Table 5.** Conservation of regions with transient α-helical structure (in orange) in Tinin's PEVK region across different species.

Helix 9964 - 9983

| | | | |
|---|---|---|---|
| A2ASS6 | 9915 | IIDVSSKAEEVKITTITRKKEVHKEKEAVYEREEAVY-EKKVHIEPW-EEPYEELETEPY | 9972 |
| A0A8I5ZUN3 | 10034 | IIDVSSKAEEVKITTITRKKEIHKEKEAVYERKEAVY-EKKVLIEPW-EEPYEELETEPY | 10091 |
| A0A8B8T085 | 8962 | IIDVSSKAEEVKITTITRKKEVQKEKEAVYEKKQAVYEEKRLFIESV-EEPYDELEMEPY | 9020 |
| A0A5F5PKC5 | 8883 | IIDVSSKAEEVKIMTITRQKEVQKEKEAVYEKKRAVYEEKKLFIESL-EEPYDELEVEPY | 8941 |
| A0A0D9RKL7 | 9949 | IIDVSSKAEEVKIMTITRKQEVQKEKEAVYEKKQAVHKEKRVFIESF-EEPYDELEVEPY | 10007 |
| A0A6P6ER96 | 9948 | IIDVSSKAEEVKITTITRKKEVQKEKEAVYEKKQAVYEEKKVYIESMVEEPYDELEVEPY | 10007 |
| A0A5F4WBB8 | 9028 | IIDVSSKAEEVKIMTITRQKEIQKEKEAMYEKKQVIHKEKKVFIESF-EEPYDELEVEPY | 9086 |
| Q8WZ42 | 9954 | IIDVSSKAEEVKIMTITRKKEVQKEKEAVYEKKQAVHKEKRVFIESF-EEPYDELEVEPY | 10012 |
| A0A2K6RWS3 | 9944 | IIDVSSKAEEVKIMTITRKQEVQKEKEAVYEKKQAVHKERRVFIESF-EEPYDELEVEPY | 10002 |
| A0A3Q7ME22 | 9976 | IIDVSSKAEEVKITTITRKKEVHKEQEAVYEKKRTVSEEKKLFIESL-EEPYDELEVEPY | 10034 |
| A0A6P6INV1 | 9028 | IIDVSSKAEEVKIKTITRKKEVQKEKEAVYEKKRAVHEEKRIFIESL-EEPYDELEVEPY | 9086 |
| A0A8M1H409 | 9025 | IVDVSSKAEEVKITTITRKKEVHKEQEAVYEKKRAVYEEKKLFIESL-EEPYDELEVEPY | 9083 |

Helix 10066 - 10071

| | | | |
|---|---|---|---|
| A2ASS6 | 10011 | QEYYEREEGYDEGEEEWEEIYHEREIIQVQKEVHE-----EL | 10046 |
| A0A8I5ZUN3 | 10130 | QEYYEREEGYDEGEEEWEEIYHEREIIQVQKEVHE-----DL | 10165 |
| A0A8B8T085 | 9059 | QEYYERAEGYDEEEEEWEETYQEGEVIQVQKEVYEGVYSLES | 9100 |
| A0A5F5PKC5 | 8980 | QEYYEREEGYDEGEEEWEETYQEREAVQVQKEVYE-----EP | 9016 |
| A0A0D9RKL7 | 10046 | QEYYEREEGYDEGEEEWEEAYQEREVIQVQKEVYE-----ES | 10082 |
| A0A6P6ER96 | 10046 | QEYYEREEGYDEGEEEWEESYQEREIIQV-QEVHE-----EL | 10080 |
| A0A5F4WBB8 | 9125 | QDYYEREEGYDEGEEEWEETYQEREVIQVQKEVYE-----ES | 9161 |
| Q8WZ42 | 10051 | QEYYEREEGYDEGEEEWEEAYQEREVIQVQKEVYE-----ES | 10087 |
| A0A2K6RWS3 | 10041 | QEYYEREEGYDEGEEEWEEAYQEREVIQVQKEVYE-----ES | 10077 |
| A0A3Q7ME22 | 10073 | QEYYEREEGYDEGEDDWEETYQEREVIQVQKEVYK-----ES | 10108 |
| A0A6P6INV1 | 9125 | QEYYEREEGYDEGEEEWEETYQEKEVIQVQKEVHE-----ES | 9160 |
| A0A8M1H409 | 9122 | QEYYEREEGYDEGEEDWEETYQEREVVQVQKEVYE-----EA | 9157 |

UniProt ID – organism name

| | |
|---|---|
| A2ASS6 | Mus musculus (Mouse) |
| A0A8I5ZUN3 | Rattus norvegicus (Rat) |
| A0A8B8T085 | Camelus ferus (Wild bactrian camel) |
| A0A5F5PKC5 | Equus caballus (Horse) |
| A0A0D9RKL7 | Chlorocebus sabaeus (Green monkey) |
| A0A6P6ER96 | Octodon degus (Degu) |
| A0A5F4WBB8 | Callithrix jacchus (White-tufted-ear marmoset) |
| Q8WZ42 | Homo sapiens (Human) |
| A0A2K6RWS3 | Rhinopithecus roxellana (Golden snub-nosed monkey) |
| A0A3Q7ME22 | Callorhinus ursinus (Northern fur seal) |
| A0A6P6INV1 | Puma concolor (Mountain lion) |
| A0A8M1H409 | Ursus maritimus (Polar bear) |

**Extended Data Table 6.** Mean values of secondary structure content (α-helix or polyproline-II helix) in selected taxa or functional groups of IDPs, calculated by DSSP in AI-IDP ensembles, are compared to mean values in all other IDPs in DisProt. Standard error of the mean, one-sided Brunner-Munzel test p-values as well as Cohen's *d* values with standard errors (estimated through 100000 bootstraps) for the intergroup comparison are reported in the table.

| Taxa/GO group | IDPs in taxa/group | | | Other IDPs | | | p-value | Cohen's *d* | DoF |
|---|---|---|---|---|---|---|---|---|---|
| | N | α-helix avg, % | α-helix SEM, % | N | α-helix avg, % | α-helix SEM, % | | | |
| Taxa: Viruses | 288 | 7.670 | 0.537 | 3101 | 6.326 | 0.123 | $1.3 \cdot 10^{-3}$ | 0.17±0.06 | 342 |
| GO: Lipid binding | 255 | 8.120 | 0.475 | 3134 | 6.304 | 0.125 | $2.9 \cdot 10^{-6}$ | 0.25±0.06 | 300 |
| GO: Virion component | 191 | 7.597 | 0.492 | 3198 | 6.798 | 6.372 | $1.5 \cdot 10^{-3}$ | 0.18±0.07 | 211 |
| | N | ppII avg, % | ppII SEM, % | N | ppII avg, % | ppII SEM, % | | | |
| GO: Animal organ morphogenesis | 214 | 12.406 | 0.494 | 3175 | 10.553 | 0.126 | $1.8 \cdot 10^{-6}$ | 0.26±0.07 | 248 |
| GO: System development | 781 | 11.847 | 0.262 | 2608 | 10.318 | 0.137 | $1.4 \cdot 10^{-9}$ | 0.21±0.04 | 1231 |
| GO: Cell communication | 1055 | 11.449 | 0.224 | 2334 | 10.319 | 0.145 | $1.0 \cdot 10^{-6}$ | 0.16±0.04 | 1900 |
| GO: Biological regulation | 2028 | 11.155 | 0.154 | 1361 | 9.948 | 0.197 | $3.8 \cdot 10^{-11}$ | 0.17±0.04 | 2956 |

**Extended Data Table 7.** Mean values of secondary structure content (α-helix or polyproline-II helix) in selected taxa of IDPs in DisProt, calculated by DSSP in AI-IDP ensembles, are compared to mean values in a different taxa selection. Standard error of the mean, one-sided Brunner-Munzel test p-values as well as Cohen's d values with standard errors (estimated through 100000 bootstraps) for the intergroup comparison are reported in the table.

| N | ppII avg, % | ppII SEM, % | | N | ppII avg, % | ppII SEM, % | p-value | Cohen's d | DoF |
|---|---|---|---|---|---|---|---|---|---|
| Archaea and Bacteria | | | vs. | Eukaryota | | | | | |
| 433 | 9.028 | 0.320 | | 2668 | 10.953 | 0.138 | $3.6 \cdot 10^{-11}$ | 0.28±0.06 | 628 |
| Non-Opisthokonta Eukaryota | | | vs. | Opisthokonta | | | | | |
| 288 | 8.874 | 0.366 | | 2380 | 11.205 | 0.148 | $1.0 \cdot 10^{-9}$ | 0.35±0.07 | 379 |
| Ascomycota | | | vs. | Eumetazoa | | | | | |
| 315 | 9.419 | 0.325 | | 2065 | 11.477 | 0.162 | $1.3 \cdot 10^{-7}$ | 0.31±0.06 | 492 |
| Protostomia | | | vs. | Deuterostomia | | | | | |
| 172 | 9.895 | 0.480 | | 1892 | 11.625 | 0.171 | $1.2 \cdot 10^{-3}$ | 0.25±0.08 | 210 |

**Extended Data Table 8.** GO terms for which associated IDP groups had strongly increased ppII helix content (p-value < $10^{-5}$). 32 of 50 terms (64 %) contain the terms regulation, signaling, development or communication. For all analyzed GO terms, only 32% contained these words.

| GO term | <ppII> with term, % | <ppII> w/o term, % | Brunner-Munzel p-value |
|---|---|---|---|
| DNA-templated transcription | 11.273 | 10.5 | 9.34E-06 |
| regulation of gene expression | 11.232 | 10.44 | 8.77E-06 |
| positive regulation of RNA metabolic process | 11.59 | 10.529 | 8.68E-06 |
| positive regulation of macromolecule metabolic process | 11.373 | 10.464 | 6.44E-06 |
| nucleic acid metabolic process | 11.075 | 10.441 | 6.07E-06 |
| regulation of developmental process | 11.677 | 10.481 | 5.25E-06 |
| nucleic acid biosynthetic process | 11.167 | 10.455 | 5.18E-06 |
| nervous system development | 11.902 | 10.459 | 3.70E-06 |
| regulation of macromolecule biosynthetic process | 11.276 | 10.413 | 3.19E-06 |
| transcription regulator activity | 11.735 | 10.545 | 2.97E-06 |
| cellular response to stimulus | 11.228 | 10.334 | 2.57E-06 |
| regulation of RNA metabolic process | 11.364 | 10.476 | 2.39E-06 |
| animal organ development | 11.808 | 10.435 | 2.30E-06 |
| signaling | 11.432 | 10.328 | 2.08E-06 |
| animal organ morphogenesis | 12.406 | 10.553 | 1.84E-06 |
| tube morphogenesis | 12.577 | 10.536 | 1.80E-06 |
| positive regulation of DNA-templated transcription | 11.739 | 10.523 | 1.53E-06 |
| positive regulation of RNA biosynthetic process | 11.739 | 10.523 | 1.53E-06 |
| cell population proliferation | 12.058 | 10.487 | 1.48E-06 |
| regulation of DNA-templated transcription | 11.43 | 10.485 | 1.18E-06 |
| cell communication | 11.449 | 10.319 | 1.00E-06 |
| regulation of RNA biosynthetic process | 11.44 | 10.482 | 9.09E-07 |
| regulation of metabolic process | 11.224 | 10.309 | 5.60E-07 |
| regulation of cell population proliferation | 12.257 | 10.488 | 5.25E-07 |
| anatomical structure morphogenesis | 11.87 | 10.426 | 3.59E-07 |
| transcription by RNA polymerase II | 11.733 | 10.48 | 3.39E-07 |
| regulation of macromolecule metabolic process | 11.307 | 10.324 | 2.89E-07 |
| positive regulation of transcription by RNA polymerase II | 12.086 | 10.518 | 2.34E-07 |
| nucleus | 11.24 | 10.236 | 2.34E-07 |
| negative regulation of cellular process | 11.344 | 10.313 | 2.24E-07 |
| chromatin | 12.42 | 10.501 | 1.74E-07 |
| tissue development | 12.077 | 10.471 | 1.51E-07 |
| anatomical structure development | 11.448 | 10.315 | 1.42E-07 |
| regulation of transcription by RNA polymerase II | 11.86 | 10.474 | 7.18E-08 |
| membrane-enclosed lumen | 11.558 | 10.277 | 7.04E-08 |
| organelle lumen | 11.558 | 10.277 | 7.04E-08 |
| intracellular organelle lumen | 11.558 | 10.277 | 7.04E-08 |
| negative regulation of biological process | 11.35 | 10.29 | 5.91E-08 |
| positive regulation of biological process | 11.38 | 10.211 | 3.42E-08 |
| tube development | 12.609 | 10.5 | 3.25E-08 |
| nucleoplasm | 11.747 | 10.338 | 2.56E-08 |
| nuclear lumen | 11.638 | 10.319 | 1.32E-08 |
| multicellular organism development | 11.617 | 10.321 | 8.84E-09 |
| developmental process | 11.503 | 10.262 | 8.50E-09 |
| positive regulation of cellular process | 11.418 | 10.226 | 5.16E-09 |
| multicellular organismal process | 11.493 | 10.23 | 4.59E-09 |
| system development | 11.847 | 10.318 | 1.39E-09 |
| regulation of cellular process | 11.183 | 10.015 | 1.28E-10 |
| biological regulation | 11.155 | 9.948 | 3.83E-11 |
| regulation of biological process | 11.181 | 9.963 | 3.38E-11 |

**Extended Data Table 9.** Comparisons of typical runtimes for different IDP ensemble generation methods and observed approximate scaling with the chain length.

| Method | 101-residue IDP, s | 236-residue IDP, s | $O(?)$ |
|---|---|---|---|
| AI-IDP | 1213 | 4302 | $O(N^2)$ |
| CALVADOS2 (+PULCHRA)* | 234 | 681 | $O(N \log N)$ |
| IDP Conf. Gen. (+MCSCE)* | 1040 | 1630000 | $\lesssim O(N^4)$ |
| IDPFold (pdbfixer+relax)** | 4371 | 15505 | $\lesssim O(N^2)$ |
| IDPFold2 (cg2all+relax)** | 2204 | 5684 | $\lesssim O(N^2)$ |
| idpGAN (cg2all+relax)** | 840 | 2231 | $O(N)$ |
| STARLING (cg2all+relax)** | 863 | 2068 | $O(N)$ |

* side chain reconstruction methods

** cg2all all-atom reconstruction + amber99db energy minimization

AI-IDP calculations were performed using Google Colab TPU v6 for AlphaFold2 fragment generation and Intel Core Ultra 9 285H for fragment assembly. CALVADOS2 calculations were performed in Google Colab using T4 GPU. IDP Conformer Generator calculations were performed on Intel Core Ultra 9 285H. IDPFold, IDPFold2, idpGAN and STARLING calculations were performed on a laptop with Intel Core Ultra 9 275HX and NVIDIA GeForce RTX 5090 GPU.
